# Degradation of Photoreceptor Outer Segments by the Retinal Pigment Epithelium Requires Pigment Epithelium-derived Factor Receptor (PEDF-R)

**DOI:** 10.1101/2021.01.01.425047

**Authors:** Jeanee Bullock, Federica Polato, Mones Abu-Asab, Alexandra Bernardo-Colón, Elma Aflaki, Martin-Paul Agbaga, S. Patricia Becerra

## Abstract

**Purpose:** To examine the contribution of PEDF-R to the phagocytosis process. Previously, we identified PEDF-R, the protein encoded by the *PNPLA2* gene, as a phospholipase A2 in the retinal pigment epithelium (RPE). During phagocytosis, RPE cells ingest abundant phospholipids and protein in the form of photoreceptor outer segment (POS) tips, which are then hydrolyzed. The role of PEDF-R in RPE phagocytosis is not known.

**Methods:** Mice in which *PNPLA2* was conditionally knocked out in the RPE were generated (cKO). Mouse RPE/choroid explants were cultured. Human ARPE-19 cells were transfected with si*PNPLA2* silencing duplexes. POS were isolated from bovine retinas. The phospholipase A2 inhibitor bromoenol lactone was used. Transmission electron microscopy, immunofluorescence, lipid labeling, pulse-chase experiments, western blots, and free fatty acid and β-hydroxybutyrate assays were performed.

**Results:** The RPE of the cKO mice accumulated lipids as well as more abundant and larger rhodopsin particles compared to littermate controls. Upon POS exposure, RPE explants from cKO mice released less β-hydroxybutyrate compared to controls. After POS ingestion during phagocytosis, rhodopsin degradation was stalled both in cells treated with bromoenol lactone and in *PNPLA2-*knocked-down cells relative to their corresponding controls. Phospholipase A2 inhibition lowered β-hydroxybutyrate release from phagocytic RPE cells. *PNPLA2* knock down also resulted in a decline in fatty acids and β-hydroxybutyrate release from phagocytic RPE cells.

**Conclusions:** PEDF-R downregulation delayed POS digestion during phagocytosis. The findings imply that efficiency of RPE phagocytosis depends on PEDF-R, thus identifying a novel contribution of this protein to POS degradation in the RPE.

---

A vital function of the retinal pigment epithelium (RPE) is to phagocytose the tips of the photoreceptors in the neural retina. As one of the most active phagocytes in the body, RPE cells ingest daily a large amount of lipids and protein in the form of photoreceptor outer segments (POS) tips.^1–5^ On the one hand, as outer segments are constantly being renewed at the base of photoreceptors, the ingestion of POS tips (~10% of an outer segment) by RPE cells serves to balance outer segment renewal, which is necessary for the visual activity of photoreceptors. On the other hand, the ingested POS supply an abundant source of fatty acids, which are substrates for fatty acid β-oxidation and ketogenesis to support the energy demands of the RPE.^6–8^ The fatty acids liberated from phagocytosed POS are also used as essential precursors for lipid and membrane synthesis, and as bioactive mediators in cell signaling processes, e.g., the main fatty acid in POS phospholipids is docosahexaenoic acid, which is involved in signaling in the retina.^9^ Rhodopsin, a pigment present in rod photoreceptors involve in visual phototransduction, is the most abundant protein in POS. Approximately 85% of the total protein of isolated bovine POS is rhodopsin,^10^ which is embedded in a phospholipid bilayer at a molar ratio between rhodopsin and phospholipids of about 1:60.^11^ Conversely, the RPE lacks expression of the rhodopsin gene. The importance of POS clearance by the RPE in the maintenance of photoreceptors was demonstrated in an animal model for retinal degeneration, the Royal College Surgeons (RCS) rats, in which a genetic defect in the RCS rats renders their RPE unable to effectively phagocytose POS, thereby leading to rapid photoreceptor degeneration.^12,13^ Moreover, human RPE phagocytosis declines moderately with age and the decline is significant in RPE of human donors with age-related macular degeneration (AMD), underscoring its importance in this disease.^14^ Therefore, there is increasing interest in studying regulatory hydrolyzing enzymes involved in RPE phagocytosis for maintaining retina function and the visual process.

We have previously reported that the human RPE expresses the *PNPLA2* gene, which encodes a 503 amino acid polypeptide that exhibits phospholipase A2 (PLA2) activity and termed pigment epithelium-derived factor receptor (PEDF-R).^15^ The enzyme liberates fatty acids from phospholipids, specifically those in which DHA is in the *sn*-2 position.^16^ RPE plasma membranes contain the PEDF-R protein,^15,17^ and photoreceptor membrane phospholipids have high content of DHA in their *sn*-2 position,^9^ suggesting that upon POS ingestion the substrate lipid is available to interact with PEDF-R. Other laboratories used different names for the PEDF-R protein (e.g., iPLA2ζ, desnutrin, adipose triglyceride lipase), and showed that it exhibits additional lipase activities: triglyceride lipase and acylglycerol transacylase enzymatic activities.^18–20^ In macrophages, the triglyceride hydrolytic activity is critical for efficient efferocytosis of bacteria and yeast.^21^ Interestingly, we and others have shown that the inhibitor of calcium-independent phospholipases A2 (iPLA2s), bromoenol lactone (BEL), inhibits the phospholipase and triolein lipase activities of PEDF-R/iPLA2ζ.^15,18^ In addition, BEL can impair the phagocytosis of POS by ARPE-19 cells, associating phospholipase A2 activity with the regulation of photoreceptor cell renewal.^22^ However, the responsible phospholipase enzyme involved in RPE phagocytosis is not yet known.

Given that the role of PEDF-R in RPE phagocytosis has not yet been studied, here we explored its contribution in this process. We hypothesized that PEDF-R is involved in the degradation of phospholipid-rich POS in RPE phagocytosis. To test this hypothesis, we silenced the *PNPLA2* gene *in vivo* and *in vitro*. Results show that with down regulation of *PNPLA2* expression and inhibition of the PLA2 activity of PEDF-R, RPE cells cannot break down rhodopsin, nor release β-hydroxybutyrate (β-HB) and fatty acids, thus identifying a novel contribution of this protein in POS degradation. We discuss the role that PEDF-R may play in the disposal of lipids from ingested OS, and in turn in the regulation of photoreceptor cell renewal.

## Methods

### Animals

The generation of desnutrin floxed mice (hereafter referred to as *Pnpla*2^f/f^)^23^ and the Tg(*BEST1-cre*)^Jdun^ transgenic line^24^ (which will be named *BEST1-cre* in this report) have been previously reported. The desnutrin floxed transgenic mouse model was kindly donated to our laboratory by Dr. Hei Sook Sul. The transgenic Tg(*BEST1-cre*)^Jdun^ mouse model was a generous gift by Dr. Joshua Dunaief. It is an RPE-specific, *cre*-expressing transgenic mouse line, in which the activity of the human *BEST1* promoter is restricted to the RPE and drives the RPE-specific expression of the targeted *cre* in the eye of transgenic mice.^24^ Homozygous floxed *Pnpla*2 (*Pnpla*2^f/f^) mice were crossed with transgenic *BEST1-cre* mice. The resulting mice carrying one floxed allele and the *cre* transgene (*Pnpla*2^f/+/cre^) were crossed with *Pnpla*2^f/f^ mice to generate mice with *Pnpla*2 knockout specifically in the RPE, which are homozygous floxed mice expressing the *cre* transgene only in the RPE, *Pnpla*2^f/f/Cre^ (here also termed cKO). *Pnpla*2^f/f/cre^ or *Pnpla*2^f/+/Cre^ were also used for breeding with *Pnpla*2^f/f^ to expand the colony. *Pnpla*2^f/+^ or *Pnpla*2^f/f^ littermates, obtained through this breeding, were used as control mice. All procedures involving mice were conducted following protocols approved by the National Eye Institute Animal Care and Use Committee and in accordance with the Association for Research in Vision and Ophthalmology Statement for the Use of Animals in Ophthalmic and Vision Research. The mice were housed in the NEI animal facility with lighting at around 280-300 lux in 12 h (6 AM-6 PM) light/12 h dark (6 PM-6 AM) cycles.

### DNA isolation

DNA was isolated from mouse eyecups using the salt-chloroform DNA extraction method^25^ and dissolved in 200 μl of TE (Tris-EDTA composed of 10 mM Tris-HCl, pH 8, and 1 mM EDTA). Aliquots (2 μl) of the DNA solution were then used for each PCR reaction using oligonucleotide primers P1 and P2 (sequences kindly provided by the laboratory of Dr. Hei Sook Sul; **Table 1**).

**Table 1.**
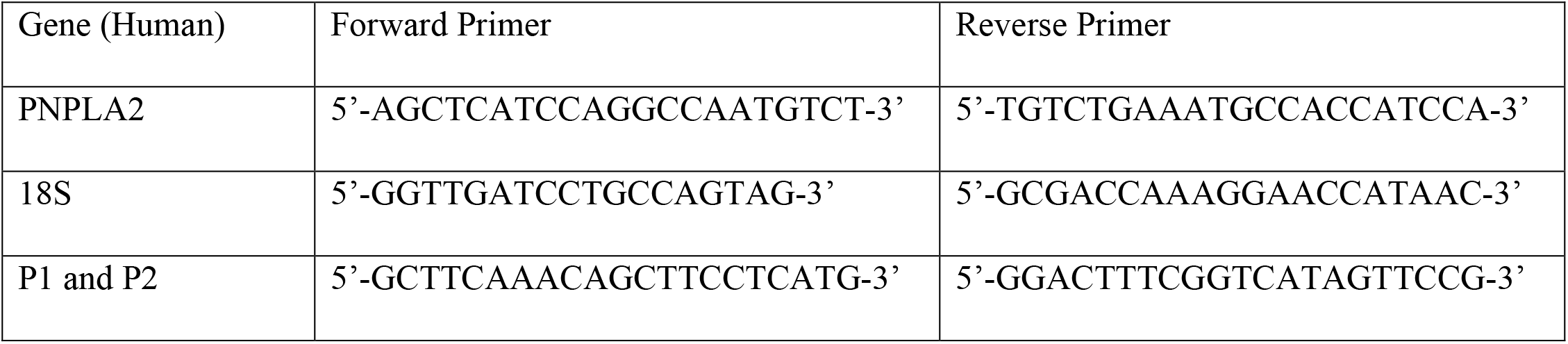
Primers used for qRT-PCR.

### RNA extraction, cDNA synthesis, and quantitative RT-PCR

RNA was isolated from the mouse RPE following the methodology previously described.^26^ Total RNA was purified from ARPE-19 cells using the RNeasy^®^ Mini Kit (Qiagen, Germantown, MD) following the manufacturer’s instructions. Between 100-500 ng of total RNA were used for reverse transcription using the SuperScript III first-strand synthesis system (Invitrogen, Carlsbad, CA). The *PNPLA*2 transcript levels in ARPE19 cells determined by quantitative RT-PCR were normalized using the QuantiTect SYBR Green PCR Kit (Qiagen) in the QuantStudio 7 Flex Real-Time PCR System (Thermo Fisher Scientific, Waltham, MA). The primer sequences used in this study are listed in **Table 1**. Murine *PNPLA*2 mRNA levels relative to *HPRT* transcript levels were measured by the QuantStudio 7 Flex Real-Time PCR System using Taqman^®^ gene expression assays (*PNPLA*2, Mm00503040_m1; *HPRT*, Mm00446968_m1, Thermo Fisher Scientific). *PNPLA*2 relative expression to *HPRT* was calculated using the comparative ΔΔCt method.^27^

### Eyecup flatmounts

Eyecup (RPE, choroid, sclera) flatmounts were prepared and processed as follows. After enucleation, and removal of cornea, lens, and retina, eyecups were fixed for 1 h in 4% paraformaldehyde at room temperature, and washed 3 times for 10 min each in Tris-Buffered Saline (TBS; 25mM Tris HCl pH 7.4, 137 mM NaCl, 2.7 mM KCl). They were then blocked for 1 h with 10% normal goat serum (NGS) in 0.1% TBS-T^a^ (TBS containing 0.1% Triton-X, Sigma, St. Louis, MO). Primary antibodies against cre recombinase and rhodopsin (see **Table 2**) in 0.1% TBS-T^a^ containing 2% NGS were diluted and used at 4°C for 16 h. Then, the eyecups were washed 3 times for 10 min each with TBS-T^a^ followed by incubation at room temperature for 1 h with the respective secondary antibodies, using DAPI (to counterstain the nuclei) and Alexa Fluor 647-phalloidin (to label the RPE cytoskeleton) diluted in 0.1% TBS-T^a^ containing 2% NGS. Eyecups were then flattened by introducing incisions and mounted with Prolong Gold antifade reagent (Thermo Fisher Scientific). Images of the entire flatmounts were collected using the tiling feature of the epifluorescent Axio Imager Z1 microscope (Carl Zeiss Microscopy, White Plains, NY) at 20X magnification. The collected images were stitched together using the corresponding feature of the Zen Blue software (Carl Zeiss Microscopy). Eyecups were also imaged using confocal microscopy (Zeiss LSM 700) at 20X magnification collecting z-stacks spanning 2 μm from each other and covering from the basal to the apical surface of the RPE cells. The image resulting from the maximum intensity projection of the z-stacks was employed for analysis.

**Table 2.** Antibodies used in the study.

| Antibody | Type & host | Application | Dilution | Company | Catalog number |
| --- | --- | --- | --- | --- | --- |
| GAPDH | Monoclonal mouse | WB | 1:10,000 | GeneTex | GTX627408 |
| PEDF-R | Polyclonal rabbit | WB<br>IF | 1:1000<br>1:250 | Protein tech | 55190-1-AP |
| Rhodopsin (A531) | Monoclonal mouse | WB<br>IF | 1:5000<br>1:800 | Novus Biologicals | NBP2-25159 |
| Rhodopsin (B630) | Monoclonal mouse | IF | 1:1000 | Novus Biologicals | NBP2-25160 |
| cre Recombinase | Monoclonal rabbit | IF | 1:800 | Cell Signaling Technology | 15036 |
| Alexa Fluor 488 | Goat anti-Mouse IgG (H+L) | IF | 1:500 | ThermoFisher Scientific | A-11001 |
| Alexa Fluor 555 | Goat anti-Rabbit IgG (H+L) | IF | 1:500 | ThermoFisher Scientific | A-21428 |
| Alexa Fluor 647 - phalloidin |  | IF | 1:100 | Cell Signaling Technology | 8940 |

Five regions of interest (ROI; 520 μm × 520 μm) were selected for each image of the flatmount from cKO mice and control mice. The percentage of cre-positive cells was determined by dividing the number of cells containing cre-stained nuclei by the number of RPE cells in each ROI (identified by F-actin staining).

For phagocytosis assay, at least six ROI (320.5 μm × 320.5 μm) were analyzed per mouse. Rhodopsin-stained particles were counted using Image J, after adjusting the color threshold and size of the particles to eliminate the background.

### Transmission electron microscopy

Mouse eyes were enucleated and doubly-fixed in 2.5% glutaraldehyde in PBS and 0.5% osmium tetroxide in PBS and embedded in epoxy resin. Thin sections (90nm in thickness) sections were generated and placed on 200-mesh copper grids, dried for 24 h, and double-stained with uranyl acetate and lead citrate. Sections were viewed and photographed with a JEOL JM-1010 transmission electron microscope.

### Electroretinography (ERG)

In dim red light, overnight dark-adapted mice were anesthetized by intraperitoneal (IP) injection of Ketamine (92.5 mg/kg) and xylazine (5.5 mg/kg). Pupils were dilated with a mixture of 1% tropicamide and 0.5 % phenylephrine. A topical anesthetic, Tetracaine (0.5%), was administered before positioning the electrodes on the cornea for recording. ERG was recorded from both eyes by the Espion E2 system with ColorDome (Diagnosys LLC, Lowell, MA, USA). Dark-adapted responses were elicited with increasing light impulses with intensity from 0.0001 to 10 candela-seconds per meter squared (sc cd.s/m2). Light-adapted responses were recorded after 2 min adaptation to a rod-saturating background (20 cd/m2) with light stimulus intensity from 0.3 to 100 sc cd.s/m2. During the recording, the mouse body temperature was maintained at 37°C by placing them on a heating pad. Amplitudes for a-wave were measured from baseline to negative peak, and b-wave amplitudes were measured from a-wave trough to b-wave peak.

### DC ERG

For DC-ERG, sliver chloride electrode connected to glass capillary tubes filled with Hank’s buffered salt solution (HBSS) were used for recording. The electrodes were kept in contact with the cornea for 10 minutes minimum until the electrical activity reached steady-state. Responses to 7-min stead light stimulation were recorded.

### Cell Culture

Human ARPE-19 cells (ATCC, Manassas, VA, USA, Cat. # CRL-2302) were maintained in Dulbecco’s modified eagle medium/Nutrient Mixture F-12 (DMEM/F-12) (Gibco; Grand Island, NY) supplemented in 10% fetal bovine serum (FBS) (Gibco) and 1% penicillin/streptomycin (Gibco) at 37°C with 5% CO_2_. For assays described below, a total of 1 × 10^5^ cells in 0.5 ml were plated per well of a 24-well plates and incubated for 3 days in DMEM/F12 with 10% FBS and 1% penicillin-streptomycin. ARPE-19 cells were authenticated by Bio-Synthesis (Lewisville, TX) at passage 27. ARPE-19 cells in passage numbers 27-32 were used for all experiments.

### Silencing of *PNPLA2* in ARPE-19 cells using siRNA

Small interfering RNA (siRNA) oligo duplexes of 27 bases in length for human *PNPLA2* were purchased from OriGene (Rockville, MD). Their sequences, and that of a Scramble siRNA (Scr) (Cat#: SR324651 and SR311349) are given in **Table 3**. From the six duplexes, siRNAs C, D, and E consistently provided the highest silencing efficiency and therefore these three duplexes were used individually for silencing experiments and referred to as si*PNPLA2*. ARPE-19 cells were transfected by reverse transfection in 24-well tissue culture plates as follows: A total of 6 pmols of siRNA was diluted in 100 μl of OptiMem (Gibco) per well, mixed with 1 μl of Lipofectamine RNAiMAX (Invitrogen), and mock transfected cells received only 1 μl of Lipofectamine. Then the mixture was added to each well. After incubation at room temperature for 10 min, a total of 1 × 10^5^ cells in 500 μl antibiotic-free DMEM/F12 containing 10% FBS was added to each well and the plate was swirled gently to mix. Assays were performed 72 h post-transfection.

**Table 3.** siRNA duplex sequences.

| siRNA Duplex | Identifier | Duplex sequences |
| --- | --- | --- |
| SR311349A | A | rCrGrCrCrArArArGrCrArCrArUrGrUrArArUrArArArUrGCT |
| SR311349B | B | rGrGrCrArCrArUrArUrArGrArArCrGrUrArCrUrGrCrArUrUCC |
| SR311349C | C | rGrCrCrUrGrArGrArCrGrCrCrUrCrCrArUrUrArCrCrArCTG |
| SR324651A | D | rCrCrArArGrUrUrCrArUrUrGrArGrGrUrArUrCrUrArArAGA |
| SR324651B | E | rCrUrGrCrCrArCrUrCrUrArUrGrArGrCrUrUrArArGrArACA |
| SR324651C | F | rCrUrUrGrGrUrArArArUrArArArArArCrGrArArArArUrGTT |

### Phagocytosis of bovine POS by ARPE-19 cells

POS were isolated as previously described^28^ from freshly obtained cow eyes (J.W. Treuth & Sons, Catonsville, MD). POS pellets were stored at −80°C until use. Quantification of POS units was performed using trypan blue and resulted in an average of 5 × 10^7^ POS units per bovine eye. The concentration of protein from purified POS was 21 pg/POS unit. Proteins in the POS samples resolved by SDS-PAGE had the expected migration pattern for both reduced and non-reduced conditions, and the main bands stained with Coomassie Blue comigrated with rhodopsin-immunoreactive proteins in western blots of POS proteins (**Fig. S1**). The percentage of rhodopsin in the protein content of POS was estimated from the gels and revealed that 80% or more of the protein content corresponded to rhodopsin.

Using electrospray ionization-mass spectrometry-mass spectrometry (ESI-MSMS) as previously described,^29^ we determined the lipid composition of the POS that were fed to the ARPE-19 cells. Phagocytosis assays in ARPE-19 cells were performed as follows: ARPE-19 cells (1 × 10^5^ cells per well) were attached to 24-well plates (commercial tissue culture-treated polystyrene plates, TCPS,^30^ purchased from Corning, Corning, NY) and cultured for 3 days to form confluent and polarized cell monolayers, as we reported previously.^31^ Ringer’s solution was prepared and composed of the following: 120.6 mM NaCl, 14.3 mM NaHCO_3_, 4.2 mM KCl, 0.3 mM MgCl_2_, and 1.1 mM CaCl_2_, with 15 mM HEPES dissolved separately and adjusted to pH 7.4 with N-methyl-D-glucamine. Prior to use, L-carnitine was added to the Ringer’s solution to achieve a 1 mM final concentration of L-carnitine. Purified POS were diluted to a concentration of 1 × 10^7^ POS/ml in Ringer’s solution containing freshly prepared 5 mM glucose. A total of 500 μl of this solution (medium) was added to each well and the cultures were incubated for 30 min, 60 min or 2.5 h, at 37°C. For pulse-chase experiments, after 2.5 h of incubation with POS (pulse), media with POS were removed from the wells and replaced with DMEM/F12 containing 10% FBS and continue incubation for a total of 16 h. The media were separated from the attached cells and stored frozen until use, and the cells were used for preparing protein extracts and either used immediately or stored frozen until used. For experiments using BEL (Sigma), BEL dissolved in vehicle dimethyl sulfoxide (DMSO) was mixed with Ringer’s solution and the mixture added to the cells and incubated for 1 h prior to starting the phagocytosis assays. The mixture was removed and replaced with the POS mixture as described above containing DMSO or BEL during the pulse. The assays were performed in duplicate wells per condition and each set of experiments were repeated at least two times.

### Cell viability by crystal violet staining

ARPE-19 cells were seeded in a 96-well plate at a density of 2 × 10^4^ cells per well. The cells were incubated at 37°C for 3 d. The medium was removed and replaced with Ringer’s solution containing various concentrations of BEL and continued incubation at 37°C for 3.5 h. The medium was replaced with complete medium and the cultures incubated for a total of 16 h. After two washes of the cells with deionized H_2_O, the plate was inverted and tapped gently to remove excess liquid. A total of 50 μl of a 0.1% crystal violet (Sigma) staining solution in 25% methanol was added to each well and incubated at room temperature for 30 min on a bench rocker with a frequency of 20 oscillations per min. The cells in the wells were briefly washed with deionized H_2_O, and then the plates were inverted and placed on a paper towel to air dry without a lid for 10 min. For crystal violet extraction, 200 μl of methanol were added to each well and the plate covered with a lid and incubated at room temperature for 20 min on a bench rocker set at 20 oscillations per min. The absorbance of the plate was measured at 570 nm.

### Western blot

ARPE-19 cells plated in multiwell cell culture dishes were washed twice with ice-cold DPBS (137 mM NaCl, 8 mM Na_2_HPO_4_-7H_2_0, 1.47 mM KH_2_PO4, 2.6 mM KCl, 490 μM MgCl_2_-6H_2_0, 900 μM CaCl_2_, pH 7.2). A total of 120 μl of cold RIPA Lysis and Extraction buffer (Thermo Fisher Scientific) with protease inhibitors (Roche, Indianapolis, IN, added as per manufacturer’s instructions) was added to each well and the plate was incubated on ice for 10 min. Cell lysates were collected, sonicated for 20 s with a 50% pulse (Fischer Scientific Sonic Dismembrator Model 100, Hampton, NH), and cellular debris are removed from soluble cell lysates by centrifugation at 20,800 × *g* at 4°C for 10 min. Protein concentration in the lysates was determined using the Pierce™ BCA Protein Assay Kit (Thermo Fisher Scientific) and the cell lysates were stored at −20°C until use. Between 5 - 10 μg of cell lysates were used for western blots.

Proteins were resolved by SDS-PAGE and transferred to nitrocellulose membranes for immunodetection. The antibodies used are listed on **Table 2**. For PEDF-R immunodetection, membranes were incubated in 1% BSA (Sigma) in TBS-T^b^ (50 mM Tris pH 7.5, 150 mM NaCl containing 0.1% Tween-20 (Sigma) at room temperature for 1 h. Then they were incubated in a solution of primary antibody against human PEDF-R at 1:1000 in 1% BSA/TBS-T^b^ at 4°C for over 16 h. Membranes were washed vigorously with TBS-T^b^ for 30 min and incubated with anti-rabbit-HRP (Kindlebio, Greenwich, CT) diluted 1:1000 in 1% BSA/TBS-T^b^ at room temperature for 30 min. The membranes were washed vigorously with TBS-T^b^ for 30 min and immunoreactive proteins were visualized using the KwikQuant imaging system (Kindlebio). For rhodopsin immunodetection, membranes were incubated in 5% dry milk (Nestle, Arlington, VA) in PBS-T (137 mM NaCl, 2.7 mM KCl, 10 mM Na_2_HPO_4_, 2 mM KH_2_PO_4_, pH 7.4, 0.1% Tween 20) at room temperature for 1 h. Then, the membranes were incubated in a solution of primary antibody against human rhodopsin (Novus, Littleton, CO) at 1:5000 in a suspension of 5% dry milk in PBS-T at 4°C for over 16 h. The membranes were washed vigorously with PBS-T for 30 min and followed with incubation in a solution of anti-mouse-HRP (Kindlebio) 1:1000 in 5% milk in PBS-T at room temperature for 30 min. The membranes were washed vigorously with PBS-T for 30 min and immunoreactive proteins were visualized using the KwikQuant imaging system. For protein loading control, the antibodies in membranes as processed described above were removed using Restore™ Western Blot Stripping Buffer (Thermo Fisher Scientific), sequentially followed by incubation with blocking 1% BSA in TBS-T at room temperature for 1 h, a solution of primary antibody against GAPDH (Genetex, cat. # GTX627408, Irvine, CA) 1:10,000 in 1% BSA/TBS-T at 4°C for over 16 h. After washing the membranes vigorously with TBS-T at room temperature for 30 min, they were incubated in a solution of anti-mouse-HRP at 1:1000 in 1% BSA/TBS-T at room temperature for 30 min. After washes with TBS-T as described above, the immunoreactive proteins were visualized using the KwikQuant imaging system.

### β-Hydroxybutyrate quantification assay

In mice, the assay was performed as described before.^8^ Briefly, after the removal of the cornea, lens and retina, optic nerve, and extra fat and muscles, the eyecup explant from one eye was placed in a well of a 96-well plate containing 170 μl Ringer’s solution and the eyecup from the contralateral eye in another well with the same volume of Ringer’s solution containing 5 mM glucose and purified bovine POS (200 μM phospholipid content, a kind gift from Dr. Kathleen Boesze-Battaglia). The eyecup explant cultures were then incubated for 2 h at 37°C with 5% CO_2_ and, the media were collected and used immediately or stored frozen until use. In ARPE-19 cells, at the endpoint of the phagocytosis assay as described above, a total of 100 μl of the culturing medium was collected and used immediately or stored at 80°C until use. The levels of β-hydroxybutyrate (β-HB) released from the RPE cells were determined in the collected samples using the enzymatic activity of β-HB dehydrogenase in a colorimetric assay from the Stanbio Beta-hydroxybutyrate LiquiColor Test (Stanbio cat. # 2440058; Boerne, TX) with β-HB standards and following manufacturer’s instructions.

### Free fatty acids quantification assay

A total of 50 μl of conditioned medium from ARPE-19 cell cultures were collected and used to quantify free fatty acids using the Free Fatty Acid Quantification Assay Kit (Colorimetric) (Abcam cat. # ab65341; Cambridge, MA) following manufacturer’s instructions.

### Statistical analyses

Data were analyzed with the two-tailed unpaired Student t test or 2-way ANOVA (analysis of variance), and are shown as the mean ± standard deviation (SD). *P* values lower than 0.05 were considered statistically significant.

## Results

### Generation of an RPE-specific *Pnpla*2-KO mouse

To circumvent the premature lethality of *PNPLA*2-KO mice,^32^ a mouse model with RPE-specific knockout of the *PNPLA*2 gene was designed. For this purpose, we crossed *Pnpla*2^f/f^ mice^23^ with *BEST1*-*cre* transgenic mice^24^ to obtain mice with conditional *Pnpla2*-knockout specific to the RPE, hereafter referred to as cKO (or *Pnpla*2^f/f/cre^). In the cKO mice, the promoter of the RPE-specific gene *VMD2* (human bestrophin, here referred as *BEST*1) drive the expression of the c*re* (*cyclization recombinase)* recombinase and restrict it to the RPE. These mice carry two floxed alleles in the *Pnpla*2 gene and a copy of the *BEST1*-c*re* transgene (*Pnpla*2^f/f/cre^).

We performed PCR reactions with primers P1 and P2, upstream and downstream from the *loxP* sites flanking exon 1, respectively (**Fig. 1A**), with DNA extracted from cKO eyecups and found that the amplimers had the expected length of 253 bp corresponding to the recombined (cKO) allele (**Fig. 1B**), thus showing that the cre-loxP recombination occurred successfully and led to the deletion of the floxed region (exon 1) in the RPE of cKO mice (or *Pnpla*2^f/f/cre^). Conversely, we observed two PCR bands of 1749 bp and 1866 bp for littermate *Pnpla*2^f/+^ control mice carrying a WT and a floxed allele, respectively (the floxed allele contains two *loxP* sites) (**Fig 1B**). In lanes for the cKO (or *Pnpla*2^f/f/cre^), we also observed very low intensity bands migrating at positions corresponding to 1749 bp and 1866 bp, which probably resulted from a few unsuccessful recombination events.

**Figure 1.**
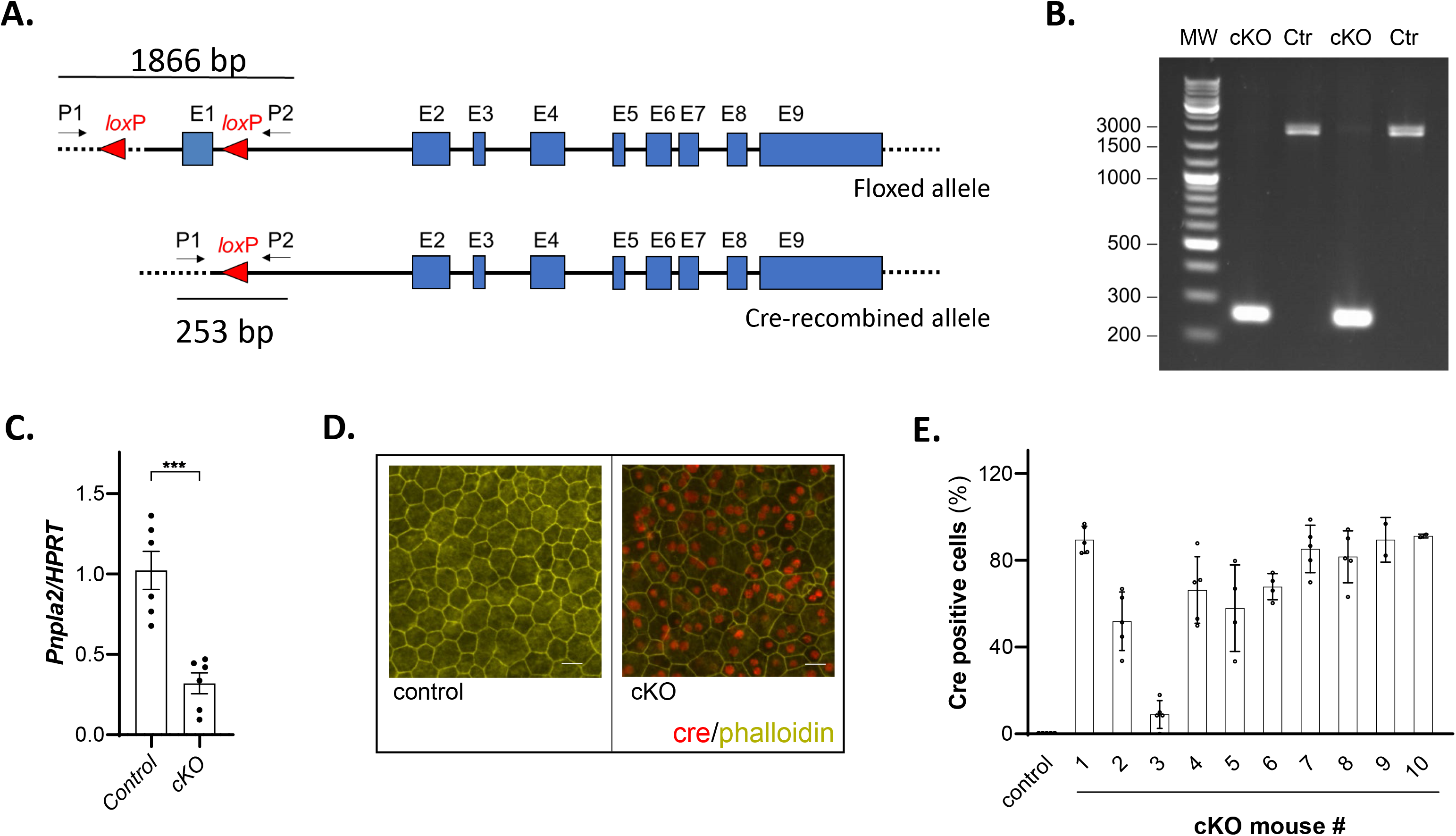
Generation of RPE-specific *PNPLA*2-cKO mice. **(A)** Scheme of *Pnpla*2 floxed and cre-mediated recombined allele. The *loxP* sites flank Exon 1. P1 and P2 are the primers homologous to sequences outside the floxed (flanked by the *loxP* sites) region used to detect cre-mediated recombination (generating recombined alleles) on genomic DNA. The sizes of the amplicons obtained by PCR using P1 and P2 are indicated. **(B)** Gel electrophoresis of PCR reaction products obtained using primers P1 and P2 and genomic DNA isolated from mouse eyecups from either cKO or control (Ctr) mice (*Pnpla*2^f/+^); lane 1 (MW) corresponds to molecular weight markers (GeneRuler DNA Ladder Mix). One eyecup per lane from a 4-month old mouse, n=2 cKO, n=2 Ctr. **(C)** *Pnpla*2 expression (*vs*. *HPRT*) in RPE from month-old cKO (*Pnpla*2^f/f/cre^) relative to control littermates (*Pnpla*2^f/f^). Each data point corresponds to the average of six PCR reactions per eyecup, six eyes from three cKO mice and six eyes from three control mice at 5 – 7 months old. **(D)** cre (red) and phalloidin (yellow) labeling of RPE/choroid flatmounts from control (*Pnpla2^f/f^*) (left) and littermate cKO *(Pnpla2^f/f/cre^*) (right). The scale corresponds to 20 μm. (n=2 images from individual mouse eyecup at 11-14 months old). **(E)** Plot of percentage of cre-positive RPE cells in cKO animals (*Pnpla2^f/f/cre^*, n=10, age was 10.5-18.5 months old) as indicated in *x-*axis. Each data point corresponds to percentage of cre-positive RPE cells from an ROI, each bar corresponds to a flatmount of an individual cKO mouse, and the bar for control (*Pnpla2^f/f^*) has data from 10 mice.

Reverse transcriptase PCR (RT-PCR) revealed *PNPLA2* transcript levels in the RPE that were lower from cKO mice than from control (with a mean that was about 32% of the control mice) (**Fig. 1C**). We determined the percentage of RPE cells that produced the cre protein by immunofluorescence of RPE whole flatmounts. Cells were visualized by co-staining with fluorescein-labelled phalloidin antibody to detect the actin cytoskeleton. We observed cre-immunoreactivity in the RPE flatmounts isolated from cKO mice, while no cre-labeling was detected in the controls (**Fig. 1D**). The overall distribution was patchy and mosaic, as previously described for the *BEST1-cre* mice.^24^ The percentage of cre-positive cells in ROI (regions of interest) of flatmounts showed nine mice with expected percentages of cre-positive cells in RPE and one with low cre-positivity (**Fig. 1E**). The average of the mean values of cre-positive cells for each cKO mouse (mouse numbers 1, 2, 4-10) was 75% (ranging between 52%-91%), which was within the expected for cre positivity in the RPE of the *BEST1-cre* mouse.^24^ Cre-positive cells were not detected in RPE of control animals (**Fig. 1D-E**). Unfortunately, further protein analysis of PEDF-R in mouse retinas was not conclusive because several commercial antibodies to PEDF-R gave high background by immunofluorescence and in western blots. Nevertheless, the results demonstrate the successful generation of RPE-specific *PNPLA*2-knock-down mice.

### Lipid accumulates in the RPE of *Pnpla2*-cKO mice

We examined the ultrastructure of the RPE by TEM imaging. Accumulation of large lipid droplets (LDs) was observed in cKO mice as early as 3 months of age compared to the control mice cohort (Fig. 2A), and LDs were still observed in the RPE of 13-month old *Pnpla*2-cKO compared to controls (**Fig. 2B**). The presence of LDs was associated with either the lack (normally seen in the basal side) (**Fig. S2A, S2H**) or the decreased thickness of the basal infoldings, and with granular cytoplasm, abnormal mitochondria (**Fig. S2B**), and disorganized localization of organelles (mitochondria and melanosomes) (**Fig. S2A**). In some cells, LDs crowded the cytoplasm and clustered together the mitochondria and melanosomes into the apical region of the cells (**Figs. S2A, S2C, S2D**); however, the number and expansion of LDs within the cells appeared to be random (**Fig. S2E**). Normal apical cytoplasmic processes were lacking; and degeneration in the outer segment (OS) tips of the photoreceptors was apparent (**Figs. S2A, S2F**). Additionally, normal phagocytosis of the OS by RPE cells was not evident, implying certain degree of impairment (**Figs. S2A, S2E, S2G**). There were apparent unhealthy nuclei with pyknotic chromatin and leakage of extranuclear DNA (enDNA), indicating the beginning of a necrotic process (**Fig. S2B**). Some RPE cells had lighter low-density cytoplasm indicating degeneration of cytoplasmic components in contrast to the denser and fuller cytoplasm in the RPE of the littermate controls (**Fig. S2I, S2J**). Thus, these observations imply that *Pnpla2* down regulation caused lipid accumulation in the RPE.

**Figure 2.**
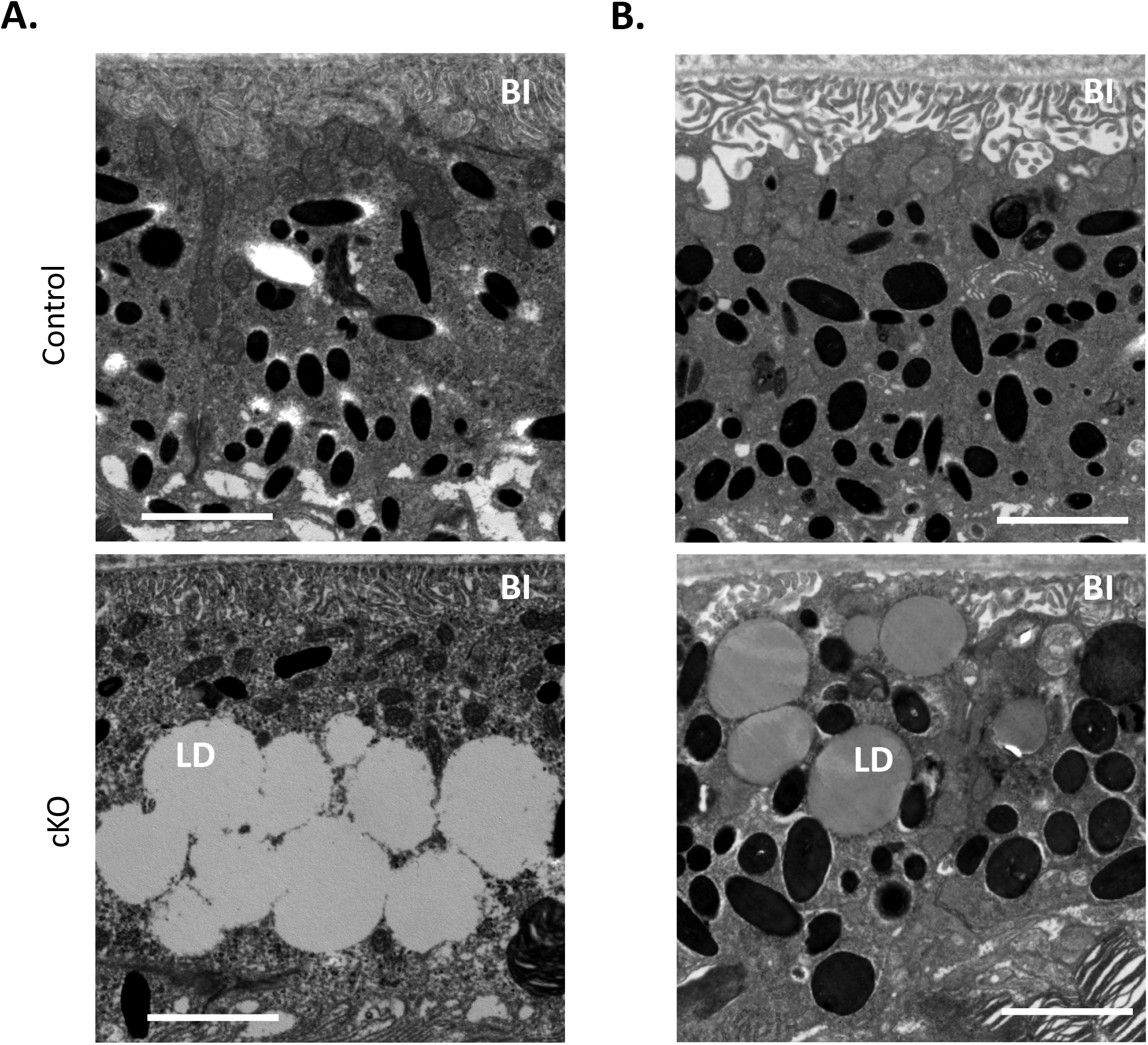
Lipid accumulation in the RPE of *Pnpla2*-cKO mice. Electron microscopy micrographs showing the RPE structure of 3- **(A)** and 13 **(B)** month-old cKO mice and control animals. LD: lipid droplets; BI: basal infoldings. Scale bar corresponds to 2 μm. The representative images were selected among examinations of micrographs from 8 eyes of cKO (*PNPLA2^f/fcre+^*) mice, from 7 eyes of control (*PNPLA2^f/f^*) mice at 1.75 - 3.75-month-old; and from 3 eyes of cKO mice and 3 eyes of control mice at 12.5 - 13-month-old.

### *Pnpla2* deficiency increases rhodopsin levels in the RPE of mice

Because the RPE does not express the rhodopsin gene, the level of rhodopsin protein in the RPE cells is directly proportional to their phagocytic activity.^5,33^ To investigate how the knock down of *Pnpla2* affects RPE phagocytic activity in mice, we compared the rhodopsin-labeled particles present in the eyecup of cKO mice and those of control mice at 2-h and 5-h post-light onset *in vivo*. The ROIs for the mutant mice were selected from areas rich in cre-positive cells. Phalloidin labeled flatmounts of control mice (n=10) showed that the RPE cells had the typical cobblestone morphology, while nine out of ten cKO mice had distorted cell morphology. Rhodopsin was detected in all ROIs and the labeled particles were more intense and larger in size in the majority of cKO flatmounts compared to those in the control mice. Representative ROIs are shown in **figure 3A**. The observations implied that *Pnpla2* knock down in the RPE prevented rhodopsin degradation *in vivo*.

**Figure 3.**
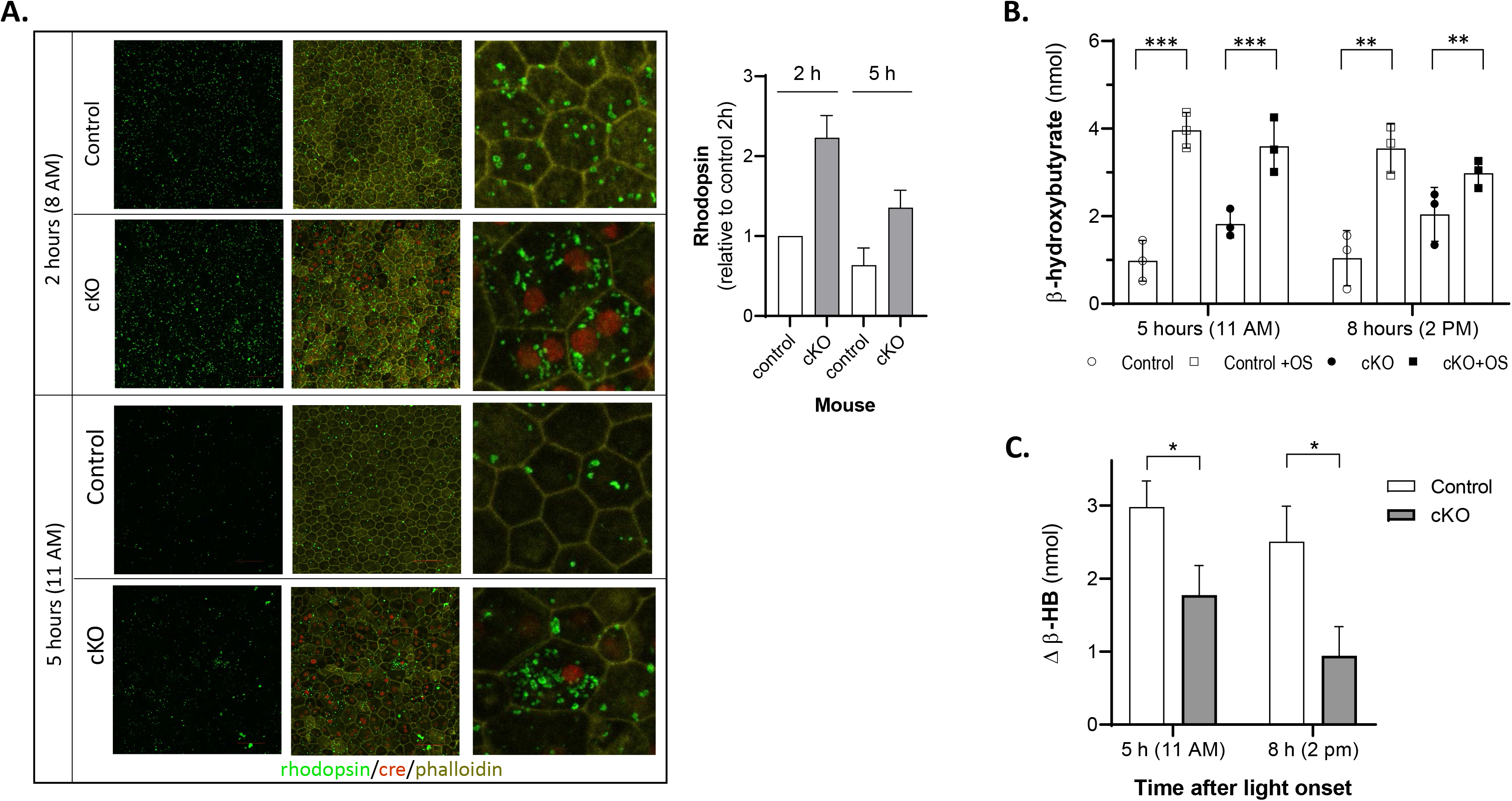
Phagocytosis and β-hydroxybutyrate production in the RPE of *Pnpla2*-cKO mice. **(A)** Representative ROI of the eyecup from one control and one cKO animal isolated at 2 h (8 AM) and 5 h (11 AM) post light onset (6 AM) after immunolabeling for rhodopsin (in green) phalloidin (in yellow) and cre (in red). The column to the right shows magnification of an area. The mean of rhodopsin immunolabel intensity in micrographs (n ≥ 6 ROIs) from flatmounts (as indicated in *x*-axis) relative to control at 2h was determined among three mice per condition and shown in the plot. Age of mice was 10.5 – 18.5 months. **(B)** *Ex-vivo* β-HB release by the RPE of *Pnpla*2-cKO eyecups upon ingestion of outer segments (OS) in comparison to that of controls. Eyecups were isolated at 5 h (11 AM) and 8 h (2 PM) after light onset (6 AM). Statistical significance was calculated using 2-way ANOVA for the 2 groups (controls and cKO mice) with and without treatment (second variance) for each time after light onset (* p=0.02; ** p=0.006; *** p=0.0001); ns, not significant. (n =6 eyecups from 3 control (f/+) mice at 3.5 months; n=4 eyecups from 2 control (f/f cre-) mice at 3.5 months; n=10 eyecups from 5 mice (f/f cre+) at 2.75 – 3.5 months) **(C)** The OS-mediated increase in β-HB release above basal levels of the cKO RPE/choroid explants was calculated from the data in Panel **(C)** and plotted.

### Ketogenesis upon RPE phagocytosis in explants from cKO mice is impaired

Given that RPE phagocytosis is linked to ketogenesis,^8^ we also measured the levels of ketone body β-HB released by RPE/choroid explants of the cKO mice *ex vivo* and compared them with those of control littermates. The experiments were performed at 5-h (11AM) and 8-h (2 PM) post-light onset, a time of day in which the amount of β-HB released due to endogenous phagocytosis is not expected to vary with time. A phagocytic challenge by exposure to exogenous bovine OS increased the amount of β-HB released by explants from both cKO and control littermates compared to the β-HB released under basal condition (without addition of exogenous OS) (**Fig. 3B**). The OS-mediated increase in β-HB release above basal levels of the cKO RPE/choroid explants (1.8 nmols at 11 AM, 0.9 nmols at 2 PM) was lower than the one of the control explants (3 nmols at 11 AM and 2.5 nmols at 2 PM) (**Fig. 3C**). These observations reveal a deficiency in β-HB production by the RPE/choroid explants of cKO mice under phagocytic challenge *ex vivo*.

### Electroretinography of the cKO mouse

To examine the functionality of the retina and RPE of cKO mice, we performed ERG and DC-ERG. Figure 4 shows histograms that revealed no differences among the animals, implying that the functionality was not affected in the RPE-*Pnpla*2-cKO mice.

**Figure 4.**
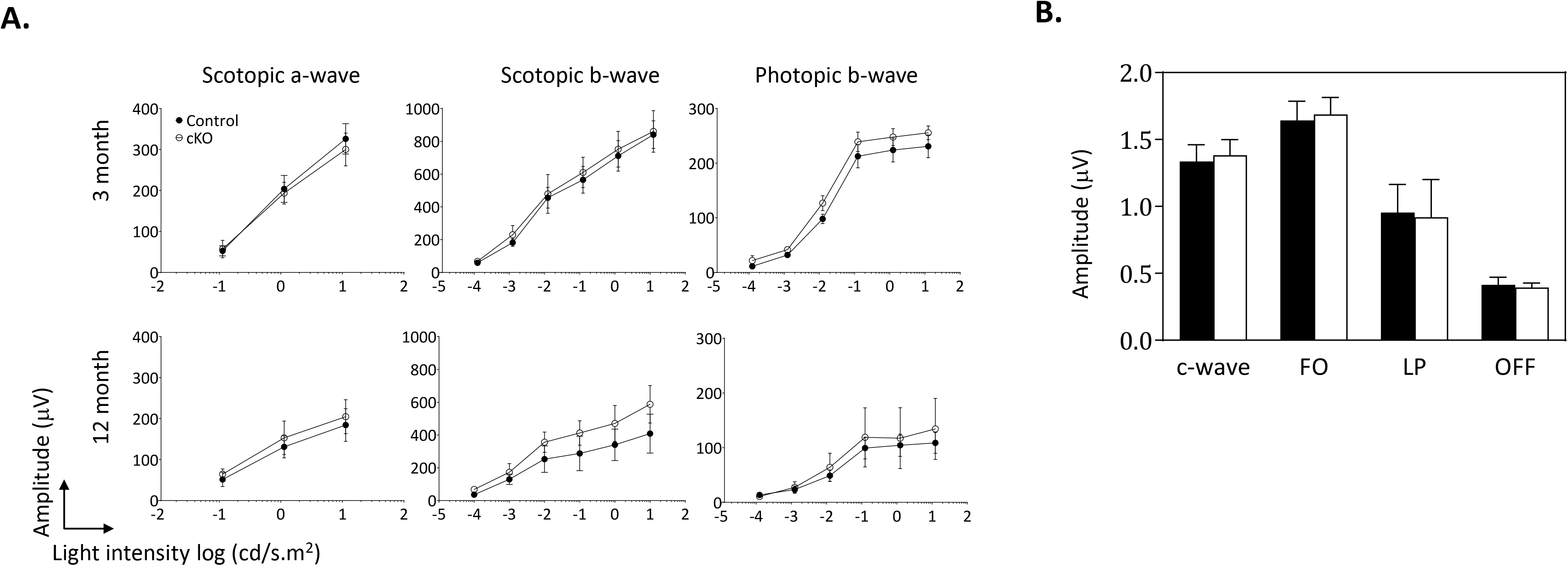
RPE and Retinal functionality in RPE-*Pnpla2-*cKO mice. (**A**) Histogram showing the amplitude (mean, standard deviation) of the c-wave, fast oscillation (FO), light peak (LP) and off-response (OFF) measured by DC-ERG of 11-week-old cKO (n=4, empty histograms) and control mice ((*Pnpla2*^f/f^ and *Pnpla2*^f/+^, n=5, filled histograms). (**B**) Electroretinograms showing amplitude (y-axis) of scotopic a- and b-wave, and photopic b-wave, as a function of light intensity (x-axis) of 3 and 12-month-old cKO mice (empty circle) and littermate controls (*Pnpla2*^f/f^, filled circles) (n=3/genotype).

### Phagocytic ARPE-19 cells engulf and break down POS protein and lipid

The complexity of the interactions that occur in the native retina makes it difficult to evaluate the subcellular and biochemical changes involved in phagocytosis of POS. Cultured RPE cells provide an ideal alternative to perform these studies. Accordingly, we designed and validated an assay with a human RPE cell line, ARPE-19, to which we added POS isolated from bovine retinas, as described in Methods. The lipid composition of the POS fed to the ARPE-19 cells included phosphatidylcholine (PC) containing very long chain polyunsaturated fatty acids (VLC-PUFAs) that was ~27 relative mole percent of total PC species in the POS. The other major PC species include PC 32:00, PC 40:06, and PC 54:10, comprising ~38 relative mole percent of the total PC phospholipids. The most abundant phosphatidylethanolamine (PE) species in the POS were PE 38:06, PE 40:05, and PE 40:06 that accounts for about 74 relative mole percent of the total PE phospholipids. The confluent monolayer of cells was exposed to the purified POS membranes for up to 2.5 h and then the ingested POS were chased for 16h for pulse-chase experiments. The fate of rhodopsin, the main protein in POS, was followed by western blotting of cell lysates. Rhodopsin was detected in the cell lysates as early as 30 min and its levels increased at 1 h and 2.5 h during the POS pulse, and decreased with a 16 h chase (**Fig. S3A**). Quantification revealed that rhodopsin levels were 21% of those detected after 2.5 h of POS supplementation (**Fig. S3B**).

Free fatty acid and β-HB levels were also determined in the culture media during the pulse. The levels of free fatty acids in the medium of POS-challenged ARPE-19 cells were 7-, 5-, and 3-fold higher at 30 min, 60 min and 2.5 h of incubation, respectively, relative to those in the medium of cells not exposed to POS (**Fig. S3C).** The β-HB levels released into the medium after POS addition also increased by 10-, 2.5- and 4-fold after 30 min, 60 min and 2.5 h incubations, respectively, relative to those observed in the medium of cells not exposed to POS (**Fig. S3D**). Altogether, these results show that under the specified conditions in this study, the batch of ARPE-19 cells phagocytosed, i.e., engulfed and digested bovine POS protein and lipid components.

### Bromoenol lactone blocks the degradation of POS components in phagocytic ARPE-19 cells

We investigated the role of PEDF-R PLA2 activity in RPE phagocytosis. As we have previously described, a calcium-independent phospholipase A2 inhibitor, bromoenol lactone (BEL), inhibits PEDF-R PLA2 enzymatic activity.^15^ First, we determined the concentrations of BEL that would maintain viability of ARPE-19 cells. **Figure 5A** shows the concentration response curve of BEL on ARPE-19 cell viability. The BEL concentration range tested was between 3.125 and 200 μM and the Hill plot estimated an IC50 (concentration that would lower cell viability by 50%) of 30.3 μM BEL. Therefore, to determine the effects of BEL on the ARPE-19 phagocytic activity, cultured cells were preincubated with the inhibitor at concentrations below the IC50 for cell viability prior to pulse-chase assays designed as described above. Pretreatment with DMSO alone without BEL was assayed as a control. Interestingly, the inhibitor at 10 μM and 25 μM blocked more than 90% of the degradation of rhodopsin during POS chase for 16 h in ARPE-19 cells **(Figs. 5B-5C)**. Similar blocking effects of BEL (25 μM) were observed with time up to 24 h during the chase (**Figs. 5D-5E**). The inhibitor did not appear to affect rhodopsin ingestion. The rhodopsin levels in pulse-chase assays with cells pretreated with DMSO alone were like those without pretreatment (compare **Figs. 5B** and **S3A**). The cells observed under the microscope after the chase point and prior to the preparation of cell lysates had similar morphology and density among cultures with and without POS, and cultures before and after pulse. Moreover, BEL blocked 40% of the β-HB releasing activity of ARPE-19 cells, whereas DMSO alone did not affect the activity **(Fig. 5F)**. These observations demonstrate that while binding and engulfment were not affected by BEL under the conditions tested, phospholipase A2 activity was required for rhodopsin degradation and β-HB release by ARPE-19 cells during phagocytosis.

**Figure 5.**
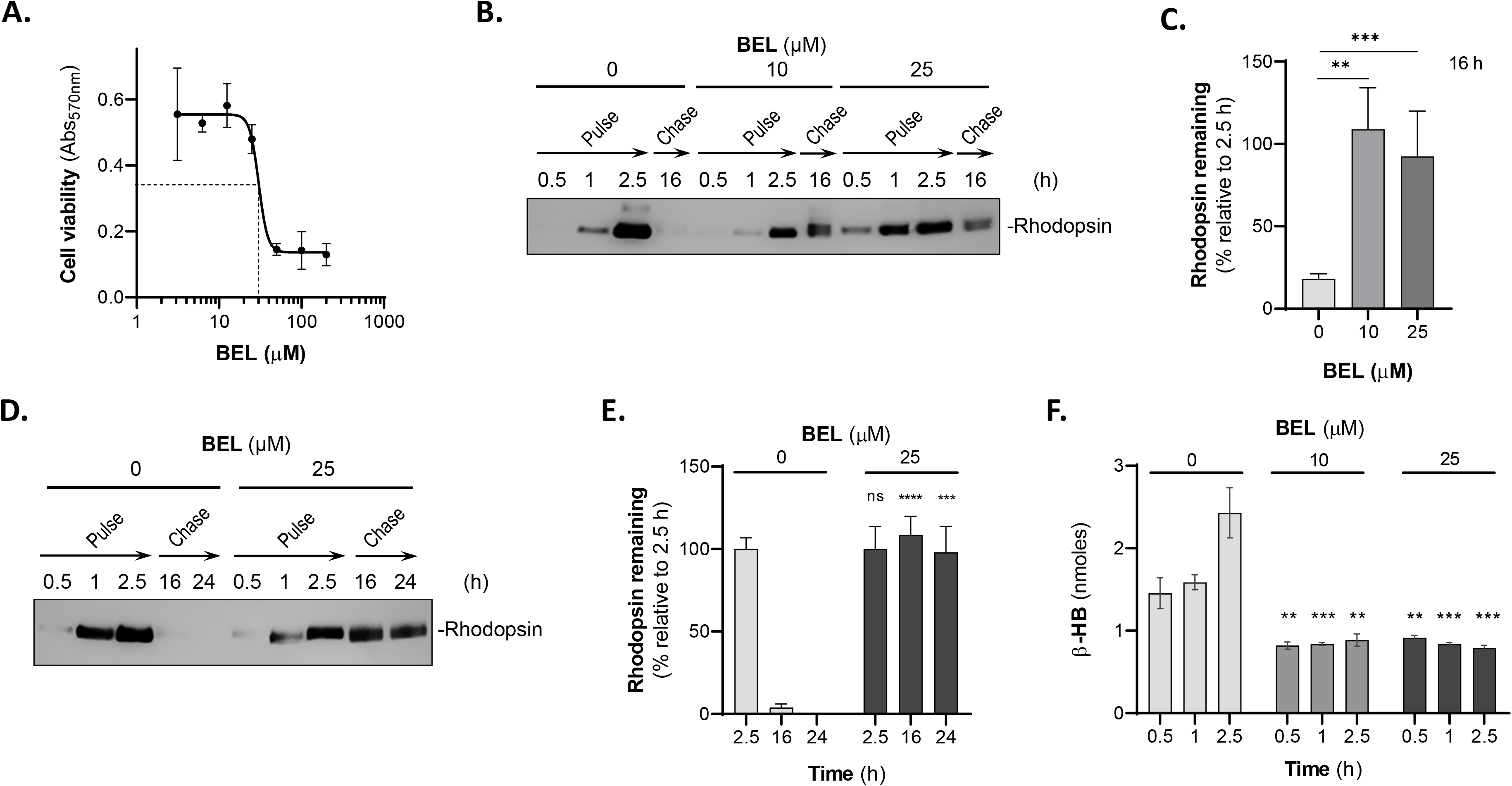
Phagocytosis in ARPE-19 cells pretreated with BEL. **(A)** ARPE-19 cells were incubated with BEL at the indicated concentrations for 3.5 h. Then the mixture was removed, washed gently with PBS, and incubated with complete medium for a total of 16 h. Cell viability was assessed by crystal violet staining and with three replicates per condition. **(B)** Representative immunoblot of total lysates of cells, which were pretreated with DMSO alone, 10 or 25 μM BEL/DMSO for 1 h prior to pulse-chase of POS, as described in methods. Extracts of cells harvested at the indicated times (*top of blot*) were resolved by SDS-PAGE followed by immunoblotting with anti-rhodopsin. Migration position of rhodopsin is indicated to the *right of the blot*. **(C)** Quantification of rhodopsin from total lysates of cells of the pulse-chase experiments as in panel **(B)**. Samples from each biological replicate were resolved in duplicate by SDS-PAGE from two experiments and single for the third experiment for quantification. Intensities of the immunoreactive bands were determined and the percentage of the remaining rhodopsin after 16-h chase relative to rhodopsin at 2.5 h-pulse was plotted. **(D)** Representative immunoblot of total lysates of cells, as in panel **B** to determine the effects of BEL at 16 h and 24 h of chase (as indicated). **(E)** Quantification of rhodopsin from two independent experiments of the pulse-chase experiments as in panel **D**. Samples from each biological replicate were resolved in duplicate by SDS-PAGE for quantification. Intensities of the immunoreactive bands were determined and the percentage of the remaining rhodopsin after 16-h chase relative to rhodopsin at 2.5 h-pulse was plotted. (**F**) Cells were preincubated with DMSO alone, 10 or 25 μM BEL/DMSO in Ringer’s solution at 37°C for 1 h. Then, the mixture was removed, and cells were incubated with Ringer’s solution containing 5 mM glucose and POS (1×10^7^ units/ml) with DMSO alone, 10 or 25 μM BEL/DMSO for the indicated times (*x*-axis). Media were removed to determine the levels of β-HB secretion, which were plotted (*y*-axis). (n=3) Data are presented as means ± S.D. **p<0.01, ***p<0.001.

### *PNPLA2* down regulation in ARPE-19 cells impairs POS degradation

We also silenced *PNPLA2* expression in ARPE-19 cells to investigate the possible requirement of PEDF-R for phagocytosis. First, we tested the silencing efficiency of six different siRNAs designed to target *PNPLA2*, along with a Scrambled siRNA sequence (Scr) as negative control (see sequences in Table 3). The siRNA-mediated knockdown of *PNPLA2* resulted in significant decreases in the levels of *PNPLA2* transcripts (siRNA A, C, D and E, **Figs. 6A** and **S5**) with a concomitant decline in PEDF-R protein levels (siRNA C, D and E, **Fig. 6D**) in ARPE-19 cell extracts. The siRNAs with the highest efficiency of silencing *PNPLA2* mRNA (namely C, D, and E) were individually used for subsequent experiments, and denoted as si*PNPLA2* (**Fig. 6A**). A time course of si*PNPLA2* transfection revealed that the gene was silenced as early as 24 h and throughout 72 h post-transfection and parallel to pulse-chase (98.5 h, **Figs. 6B, S5**). There was no significant difference between mock transfected cells and cells transfected with Scr (**Fig. 6C**). Examining the cell morphology under the microscope, we did not notice differences between the scrambled and *siPNPLA2*-transfected cells. Western blots showed that protein levels of PEDF-R in ARPE-19 membrane extracts declined 72 h post-transfection (**Fig. 6D**). Thus, subsequent experiments with cells in which *PNPLA2* was silenced were performed 72 h after transfection. Second, we tested the effects of *PNPLA2* silencing on ARPE-19 cell phagocytosis. Here we monitored the outcome of rhodopsin in pulse-chase experiments. Interestingly, while *PNPLA2* knock down did not affect ingestion, the si*PNPLA2*-transfected cells failed to degrade the ingested POS rhodopsin (88% and 24% remaining at 16 h and at 24h, respectively), while Scr-transfected cells were more efficient in degrading them (21% and 12% remaining at 16 and 24 h respectively) (**Figs. 7A-7B**).

**Figure 6.**
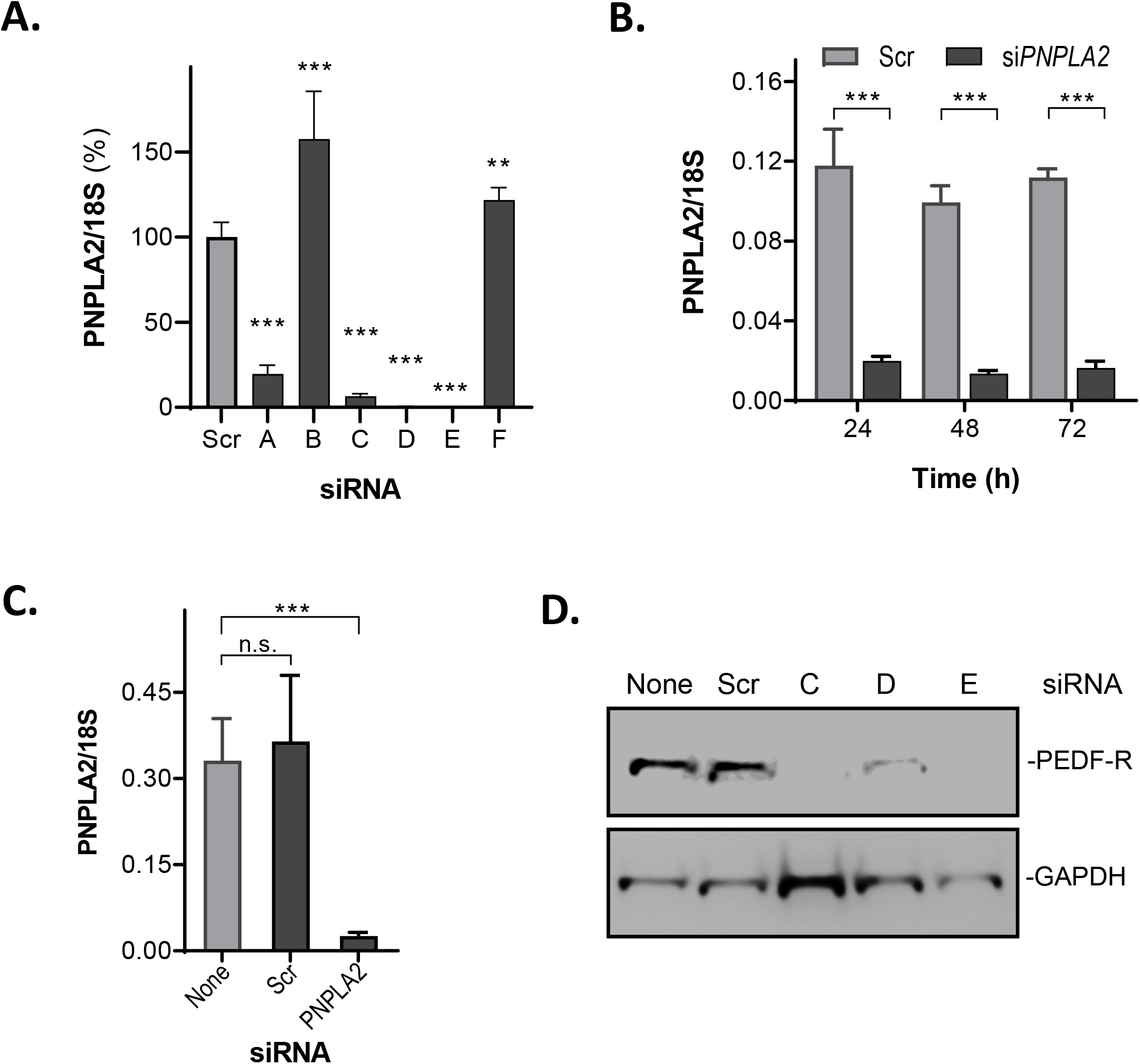
Knockdown of *PNPLA2* in ARPE-19 cells. ARPE-19 cells were transfected with Scr (Scrambled siRNA control) or siRNAs targeting *PNPLA2*, and mRNA levels and protein were tested. **(A)** RT-qPCR to measure *PNPLA2* mRNA levels in ARPE-19 cells 72 h post-transfection with Scr and six different siRNAs (as indicated in the *x*-axis) was performed and a plot is shown. *PNPLA2* mRNA levels were normalized to 18S. All siRNA are represented as the percentage of the scrambled siRNA control. n = 3 **(B)** A plot is shown for a time course of *PNPLA2* mRNA levels following transfection with Scr and *siPNPLA2-*C. n = 3 (C) RT-qPCR of mock-transfected cells, cells transfected with Scr, and *siPNPLA2-C* (*x*-axis) at 72 h after transfection. mRNA levels were normalized to the 18S RNA (*y*-axis). n = 3 **(D)** Total protein was obtained from cells harvested 72 h after transfection and resolved by SDS-PAGE followed by western blotting with anti-PNPLA2 and anti-GAPDH (loading control). The siRNAs used in transfections are indicated at the top, and migration positions for PEDF-R and GAPDH are to the right of the blot. Data are presented as means ± S.D. **p<0.01, ***p<0.001***p<0.001

**Figure 7.**
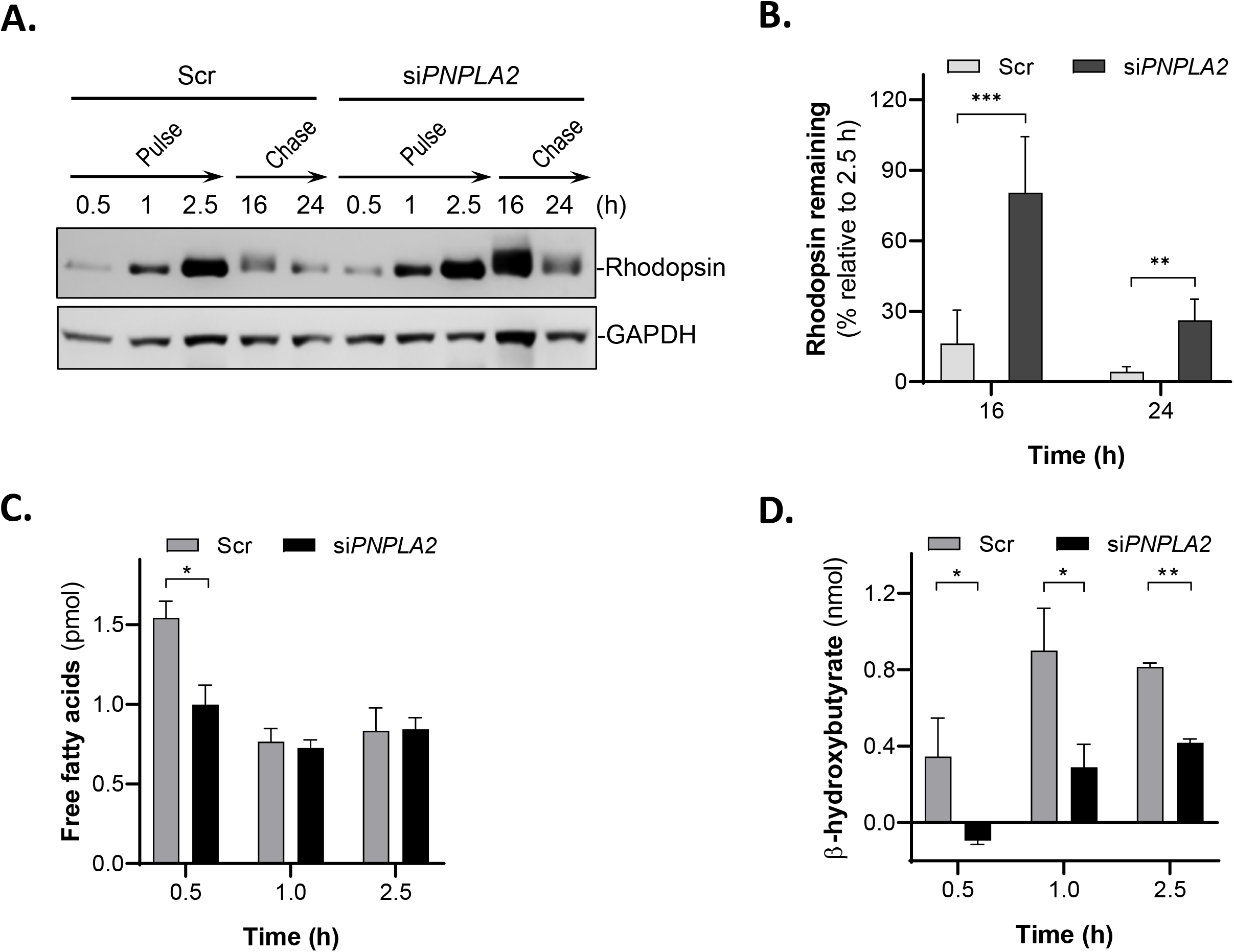
Phagocytosis and fatty acid metabolism in si*PNPLA2* cells. ARPE-19 cells were transfected with Scr or siRNAs targeting *PNPLA2.* At 72 h post-transfection, ARPE-19 cells were incubated with POS (1 × 10^7^ units/ml) in 24-well tissue culture plates for pulse-chase experiments. **(A)** Representative immunoblot of total lysates of ARPE-19 cells at 0.5 h, 1 h, and 2.5 h of POS pulse and at a 16-h and 24-h chase period, as indicated at the top of the blot. Proteins in cell lysates were subjected to immunoblotting with anti-rhodopsin followed by reprobing with anti-GAPDH as the loading control. **(B)** Quantification of rhodopsin from duplicate samples and 3 blots of cell lysates from pulse-chase experiments and time periods (indicated in the *x*-axis) as from panel. Data are presented as means ± S.D. ns, not significant, **p<0.01. **(A)**. Intensities of the immunoreactive bands were determined and the percentage of the remaining rhodopsin after 16-h and 24-h chase relative to rhodopsin at 2.5 h-pulse was plotted (*y*-axis). **(C-D)** Levels of secreted free fatty acids **(C)** and β-HB **(D)** were measured in culture media of cells transfected with Scr or *siPNPLA2* following incubation with POS for the indicated periods of times (x-axis). (n =3) Data are presented as means ± S.D. * p < 0.05, **p<0.01. Duplex si*PNPLA2* C was used to generate the data (see Table 3 for sequences of duplexes).

Third, we also determined the levels of secreted free fatty acids and β-HB production in *PNPLA2* silenced cells at 0.5 h, 1 h, and 2.5 h following POS addition. Free fatty acid levels in the culture medium were lower in *siPNPLA2*-transfected cells than in cells transfected with Scr at 30 min post-addition of POS, and no difference was observed between *siPNPLA2* and Scr at 1 h and 2.5 h post-addition (**Fig. 7C**). Secreted β-HB levels in the culture medium were lower in si*PNPLA2* cells than in Scr-transfected cells at all time points (**Fig. 7D**). To determine the effect of *PNPLA2* knockdown on lipid and fatty acid levels in the ARPE-19 cells fed POS membranes, we used electron spray ionization-mass spectrometry (ESI/MS/MS) and gas chromatography-flame ion detection to identify and quantify total lipids and fatty acid composition of the ARPE-19 cells at 2.5 and 16 h post POS feeding. Our results did not show any significant differences in the intracellular lipid and fatty acid levels in the *siPNPLA2* knockdown in Scr and WT control cells at both 2.5 and 16 h after POS addition (data not shown). Taken together, these results demonstrate that digestion of POS protein and lipid components was impaired in *PNPLA2* silenced ARPE-19 cells undergoing phagocytosis.

## Discussion

Here, we report that PEDF-R is required for efficient degradation of POS by RPE cells after engulfment during phagocytosis. This conclusion is supported by the observed decrease in rhodopsin degradation, in fatty acid release and in β-HB production upon POS challenge when the *PNPLA2* gene is downregulated or the PEDF-R lipase is inhibited. These observations occur in RPE cells *in vivo*, *ex vivo* and *in vitro*. The findings imply that RPE phagocytosis depends on PEDF-R for the release of fatty acids from POS phospholipids to facilitate POS protein hydrolysis, thus identifying a novel contribution of this enzyme in POS degradation and, in turn, in the regulation of photoreceptor cell renewal.

This is the first time that the *PNPLA2* gene has been studied in the context of RPE phagocytosis of POS. Previously, we investigated its gene product, termed PEDF-R, as a phospholipase-linked cell membrane receptor for pigment epithelium-derived factor (PEDF), a retinoprotective factor encoded by the *SERPINF1* gene and produced by RPE cells.^15,17,34,35^ Like RPE cells, non-inflammatory macrophages are phagocytic cells, but unlike RPE cells, they are found in all tissues, where they engulf and digest cellular debris, foreign substances, bacteria, other microbes, etc.^36,37^ The Kratky laboratory reported data on the effects of *PNPLA2* silencing in efferocytosis obtained using *PNPLA2*-deficient mice (termed *atgl^−/−^* mouse), and demonstrated that their macrophages have lower triglyceride hydrolase activity, higher triglyceride content, lipid droplet accumulation, and impaired phagocytosis of bacterial and yeast particles,^21^ and that in these cells, intracellular lipid accumulation triggers apoptotic responses and mitochondrial dysfunction.^38^ We have shown that *PNPLA2* gene knockdown causes RPE cells to be more responsive to oxidative stress-induced death.^39^ *PNPLA2* gene silencing, PEDF-R peptides blocking ligand binding, and enzyme inhibitors abolish the activation of mitochondrial survival pathways by PEDF in photoreceptors and other retinal cells.^17,34,40^ Consistently, overexpression of the *PNPLA2* gene or exogenous additions of a PEDF-R peptide decreases both the death of RPE cells undergoing oxidative stress and the accumulation of biologically detrimental leukotriene LTB4 levels.^31^ The fact that PEDF is a ligand that enhances PEDF-R enzymatic activity, suggests that exposure of RPE to this factor is likely to enhance phagocytosis. These implications are unknown and need further study. Exogenous additions of recombinant PEDF protein to ARPE-19 cells undergoing phagocytosis did not provide evidence for such enhancement (JB personal observations). This suggests that heterologous *SERPINF1* overexpression in cells and/or an animal model of inducible knock-in of *Serpinf1* may be useful to focus on the role of PEDF/PEDF-R in RPE phagocytosis unbiased by the endogenous presence of PEDF.

To investigate the consequences of *PNPLA2* silencing in POS phagocytosis, we generated a mouse model with a targeted deletion of *Pnpla2* in RPE cells in combination with the *BEST-cre* system for its exclusive conditional silencing in RPE cells (cKO mouse). These mice are viable with no apparent changes in other organs and in weight compared with control littermates and wild type mice. The cKO mice live to an advanced age, in contrast to the constitutively silenced *PNPLA2*-KO mice in which the lack of the gene causes premature lethality (12-16 weeks) due to heart failure associated with massive accumulation of lipids in cardiomyocytes.^32^ The RPE cells of the cKO mouse have large lipid droplets at early and late age (**Figs. 2A, S2**) consistent with a buildup of substrates for the lipase activities of the missing enzyme. In cKO mice, lipid accumulation associates with lack of or the decreased thickness of the basal infoldings, granular cytoplasm, abnormal mitochondria and disorganized localization of organelles (mitochondria and melanosomes) in some RPE cells (**Fig. S2**). Taken together, the TEM observations in combination with the greater rhodopsin accumulation and decline in β-HB release in cKO mice support that PEDF-R is required for lipid metabolism and phagocytosis in the RPE. However, interestingly, the observed features do not seem to affect photoreceptor functionality (**Fig. S3**) and appear to be inconsequential to age-related retinopathies in the *Pnpla2*-cKO mouse. This unanticipated observation suggests that the remaining RPE cells expressing *Pnpla2* gene probably complement activities of those lacking the gene, thereby lessening photoreceptor degeneration and dysfunction in the cKO mouse. We note that the cKO mouse has a mosaic expression pattern with non-cre-expressing RPE cells, as shown before for the *BEST1-cre* transgenic line.^24^ At the same time, the ERG measurements performed correspond to global responses of the photoreceptors and RPE cells, thereby missing individual cell evaluation. The lack of photoreceptor dysfunction with RPE lipid accumulation due to *PNPLA2* down regulation also suggests that during development a compensatory mechanism independent of *Pnpla2*/PEDF-R is likely to be activated, thereby minimizing retinal degeneration in the cKO mouse. Further study will be required to understand the implications of these unexpected findings. Animal models of constitutive heterozygous knockout or inducible knockdown of *PNPLA2* may be instrumental to address the role of *PNPLA2*/PEDF-R in mature photoreceptors unbiased by compensatory mechanisms due to low silencing efficiency or during development.

Results obtained from experiments using RPE cell cultures further establish that PEDF-R deficiency affects phagocytosis. It is worth mentioning that the data obtained under our experimental conditions were essentially identical to those typically obtained in assays performed with cells attached to porous permeable membranes, and this provides an additional advantage to the field by requiring shorter time to complete (see **Fig. S4**). On one hand, the decrease in the levels of β-HB and in the release of fatty acids (the breakdown products of phospholipids and triglycerides) upon POS ingestion by cells pretreated with BEL as well as transfected with si*PNPLA2* relative to the control cells indicates that *PNPLA2* participates in RPE lipid metabolism. On the other hand, the fact that PEDF-R inhibition and *PNPLA2* down regulation impair rhodopsin break down from ingested POS in RPE cells implies a likely dependence of PEDF-R-mediated phospholipid hydrolysis for POS protein proteolysis. In this regard, we envision that proteins in POS are mainly resistant to proteolytic hydrolysis, because the surrounded phospholipids block their access to proteases for cleavage. Phospholipase A2 activity would hydrolyze these phospholipids to likely liberating the proteins from the phospholipid membranes and become available to proteases, such as cathepsin D, an aspartic protease responsible for 80% of rhodopsin degradation.^41^ It is important to note that the findings cannot discern whether PEDF-R is directly associated to the molecular pathway of rhodopsin degradation, or indirectly involved in downregulating cathepsin D or other proteases. It is also possible that *PNPLA2* deficiency results in the alteration of critical genes regulating the phagocytosis pathway, such as LC3 and genes of the mTOR pathway. Animal models deficient in such genes display retinal phenotypes such as impaired phagocytosis and lipid accumulation, similar to those observed in PEDF-R deficient cells.^42–44^ These implications need further exploration.

Given that BEL is an irreversible inhibitor of iPLA2 it has been used to discern the involvement of iPLA2 in biological processes. Previously, we demonstrated that BEL at 1 to 25 μM blocks 20 – 40% of the PLA activity of human recombinant PEDF-R.^15^ Jenkins et al showed that 2 μM BEL inhibits >90% of the triolein lipase activity of human recombinant PEDF-R (termed by this group as iPLA2ζ).^18^ In cell-based assays, Wagner et al showed BEL at 20 μM inhibits 40% of this enzyme’s triglyceride lipase activity in hepatic cells.^45^ In the present study, to minimize cytotoxicity and ensure inhibition of the iPLA2 activity of PEDF-R in ARPE-19 cells, we selected 10 μM and 25 μM BEL concentrations that are below the IC50 determined for ARPE-19 cell viability (30.2 μM BEL; Fig. 5A). We note that these BEL concentrations are within the range used in an earlier study on ARPE-19 cell phagocytosis.^22^ We compared our results to those by Kolko et al ^22^ regarding BEL effects on phagocytosis of ARPE-19 cells. Using Alexa-red labeled-POS, they reported the percent of phagocytosis inhibition caused by 5 – 20 μM BEL as 24% in ARPE-19 cells. However, the authors did not specify the time of incubation for this experiment and, based on the other experiments in the report, the time period may have lasted at least 12 h of pulse, implying inhibition of ingestion of POS, and lacking description of the effects of BEL on POS degradation. With unmodified POS in pulse-chase assays, our findings show a percent of inhibition after chase of >90% for 10 μM and 25 μM BEL, indicating more effective inhibition of POS digestion. The effect of BEL on POS ingestion under 2.5 h was insignificant and over 2.5 h remains unknown (pulse). In addition, we show that pretreatment with BEL results in a decrease in the release of β-HB, which is produced from the oxidation of fatty acids liberated from POS. Thus, our assay provides new information -e.g., pulse-chase, use of unmodified POS, β-HB release- to those reported by Kolko et al. It is concluded that BEL can impair phagocytic processes in ARPE-19 cells. While BEL is recognized as a potent inhibitor of iPLA2, it can also inhibit non-PLA2 enzymes, such as magnesium-dependent phosphatidate phosphohydrolase and chymotrypsin.^46,47^ Consequently, a complementary genetic approach targeting PEDF-R is deemed reasonable and appropriate to investigate its role in RPE phagocytosis. The complex and highly regulated phagocytic function of the RPE also serves to protect the retina against lipotoxicity. By engulfing lipid-rich POS and using ingested fatty acids for energy, the RPE prevents the accumulation of lipids in the retina, particularly phospholipids, which could trigger cytotoxicity when peroxidized.^48,49^ In this regard, the lack of observed differences in intracellular phospholipid and fatty acids between PEDF-R-deficient RPE and control cells lead us to speculate that in ARPE-19 cells exposed to POS the undigested lipids remain within the cells and contribute to the total lipid and fatty acid pool, some of which may be converted to other lipid byproducts to protect against lipotoxicity. Also, the duration of the *in vitro* chase is shorter than what pertains *in vivo*, where undigested POS accumulate and overtime coalesce to form the large lipid droplets observed in the RPE *in vivo*. Thus, future experiments aimed at detailed time-dependent characterization of specific lipid species and free fatty acid levels in the RPE *in vivo*, and in media and cells *in vitro* will allow us to have a better understanding of classes of lipids and fatty acids that contribute to the lipid droplet accumulation in the RPE *in vivo* due to *PNPLA2* deletion. Nonetheless, a role of PEDF-R in POS degradation agrees with the previously reported involvement of a phospholipase A2 activity in the RPE phagocytosis of POS^22^, and with the role of providing protection of photoreceptors against lipotoxicity.

In conclusion, this is the first study to identify a role for PEDF-R in RPE phagocytosis. The findings imply that efficient RPE phagocytosis of POS requires PEDF-R, thus highlighting a novel contribution of this protein in POS degradation and its consequences in the regulation of photoreceptor cell renewal.

## Abbreviations

AMD: age-related macular degeneration
BEL: bromoenol lactone
β-HB: beta hydroxybutyrate
cre: cyclization recombinase
DHA: docosahexaenoic acid
*loxP*: locus of X-over, P1
PEDF-R: pigment epithelium-derived factor receptor
PNPLA2: patatin-like phospholipase domain containing 2
POS: photoreceptor outer segments
ROI: regions of interest
RPE: retinal pigment epithelium
TEM: transmission electron microscopy
WT: wild type

## Acknowledgements

This work was supported by the Intramural Research Program of the National Eye Institute, NIH (Project #EY000306) to SPB and by NIH/NEI R01 EY030513 to MPA. We thank the NEI animal house, Histopathology, Visual Function, Genetic Engineering and Biological Imaging Core facilities for technical support. We thank Dr. Hei Sook Sul, University of California, Berkeley, for kindly providing sequences for primers of *Pnpla2* and the Desnutrin flox mouse; Dr. Joshua Dunaief, University of Pennsylvania for kindly providing the transgenic Tg(*BEST1-cre*)^Jdun^ mouse model; Dr. Kathleen Boesze-Battaglia’s laboratory for kindly providing POS; Drs. Eugenia Poliakov and Sheetal Uppal for help in isolating POS; Dr. Kiyoharu J Miyagishima for performing the dcERG experiments; Dr. Preeti Subramanian for technical assistance with cell culture and microscopy; and Dr. Ivan Rebustini for proofreading the manuscript and providing feedback and reagents for RT-PCR.

## Supplementary Information

**Figure S1**. Proteins in the POS samples were determined and resolved by SDS-PAGE in the same gel in two sets: one with 5 μg and another with 0.1 μg protein per lane. For each set, one sample was non-reduced and the other was reduced with DTT. After electrophoresis, the gels were cut in half lengthwise. The gel portion with 5 μg of protein was stained with Coomassie Blue and the other portion with 0.1 μg protein was transferred to a nitrocellulose membrane for immunostaining using anti-rhodopsin antibodies (as described in Methods). Photos of the stained gel and western blot are shown.

The proteins of POS isolated from bovine retina had the expected migration pattern for both reduced and non-reduced conditions, and the main bands stained with Coomassie Blue comigrated with rhodopsin-immunoreactive proteins in western blots of POS proteins.

**Figure S2. Electron microscopy micrographs.** Panels **A-J** show electron microscopy micrographs of RPE structures of 3-month-old RPE cKO prepared as described in the main text and Figure 2. Magnification is indicated for each image.

The presence of LDs was associated with lack (**Fig. S2A**) of or the decreased thickness of the basal infoldings, and with granular cytoplasm, abnormal mitochondria (**Fig. S2B**), and disorganized localization of organelles (mitochondria and melanosomes) (**Fig. S2A**). In some cells, the large LDs crowded the cytoplasm and clustered together the mitochondria and melanosomes into the apical region of the cells (**Figs. S2A, S2C, S2D**); however, LDs number and expansion within the cells appeared to be random and their expansion could go into any direction (**Fig. S2E**). Normal apical cytoplasmic processes were lacking; however, degeneration in the outer segment (OS) tips of the photoreceptors was visible (**Figs. S2A, S2F**). Additionally, normal phagocytosis of the OS was lacking indicating an impaired RPE phagocytosis (**Figs. S2A, S2E, S2G**). There were apparent unhealthy nuclei with pyknotic chromatin and leakage of extranuclear DNA (enDNA), indicating that the beginning of the necrotic process had started (**Fig. S2B**). Some RPE cells lacked basal infoldings, normally seen at the basal side (**Fig. S2H**). Occasionally some RPE cells had lighter low-density cytoplasm indicating degeneration of cytoplasmic components in contrast to the denser and fuller cytoplasm in the RPE of the littermate control (**Fig. S2I, S2J**).

**Figure S3.**

**Phagocytosis in ARPE-19 cells**. ARPE-19 cells were cultured in 24-well plates for 3 days, and then exposed to POS at 1×10^7^ units/ml for up to a 2.5-h pulse followed by a 16-h chase period as described in Methods. **(A)** Representative immunoblots of total cell lysates during pulse-chase (times indicated at the top of the blot) with anti-rhodopsin followed by reprobing with anti-GAPDH as the loading control are shown. Migration positions of rhodopsin and GAPDH are indicated to the right of the blot. Duplicate biological replicates were performed. **(B)** Quantification of rhodopsin from duplicate samples per condition from pulse-chase experiments at time periods indicated in the x-axis as from panel **(A)**. Intensities of the immunoreactive bands from duplicate samples of cell lysates were determined. The percentage of the remaining rhodopsin after 16-h chase relative to rhodopsin at 2.5 h-pulse was plotted. **(C-D)** Levels of free fatty acids **(C)** and β-HB **(D)** measured in culture media of cells incubated with and without POS for the indicated periods of time (*x*-axis) were plotted and shown. n = 3 Data are presented as means ± S.D. * p < 0.05, ***p<0.001.

**Figure S4. Phagocytosis in ARPE-19 cells in porous membranes**. ARPE-19 cells were treated with 1×10^7^ POS/ml. **(A)** Representative immunoblot showing rhodopsin internalization from total cell lysates of ARPE-19 cells following 30, 60, and 150 min of POS incubation following plating in 12-well transwell inserts for 3 weeks. Cell extracts were resolved by SDS-PAGE followed by immunoblotting with anti-rhodopsin. The blot was stripped and reprobed with anti-GAPDH as a loading control. **(B)** Levels of B-HB secreted towards the apical membrane of ARPE-19 cells following POS incubation for 30, 60, and 150 min. (n = 3) Data are presented as means ± S.D.

## Methods

To demonstrate a functional assay to study phagocytosis in ARPE-19 cells we perform the assay with confluent cells attached on porous membranes

ARPE-19 cells seeded on porous membranes were incubated for 3 weeks in culturing media. Then the media was removed and replaced with Ringer’s solution alone or Ringer’s solution containing 1 × 10^7^ POS/ml and 5 mM glucose for the indicated time points. Rhodopsin was detected by western blotting.

Rhodopsin levels in the lysates of cells incubated with POS were detected in as little as 30 min and up to 2.5 h following POS incubation, while rhodopsin was undetectable in cells without POS (**Fig. S4A**). β-HB levels released into the media of the apical chamber of transwells following POS incubation increased four-fold and three-fold after 1 h and 2.5 h, respectively, while released β-HB levels from cells incubated with Ringer’s solution alone did not increase (**Fig. S4B**).

**Figure S5:** ARPE-19 cells were transfected with siScramble siRNA control or siRNAs targeting *PNPLA2 (siPNPLA2 A)*. RT-qPCR to measure *PNPLA2* mRNA levels in ARPE-19 cells at **(A)** 72 h post-transfection and (**B**) 98.5h post transfection equivalent to pulse (2.5h) and chase (24h) was performed with siRNA duplexes (as indicated in the *x*-axis). Treatment of cells in panel B was as for pulse-chase (see diagram in Fig S3). *PNPLA2* mRNA levels were normalized to 18S. n =3 biological replicates, each data point corresponds to the average of triplicate PCR reactions. The RT-PCR was repeated twice per biological replicate. Values that fell out of the standard curve were not included in the plot.

The data shows that si*PNPLA2* duplex silenced *PNPLA2* in ARPE-19 at 72 h post-transfection and that silencing was maintained throughout a 2.5 h and pulse-chase of 24 h.

## Supplementary Figures

**Figure S1.**
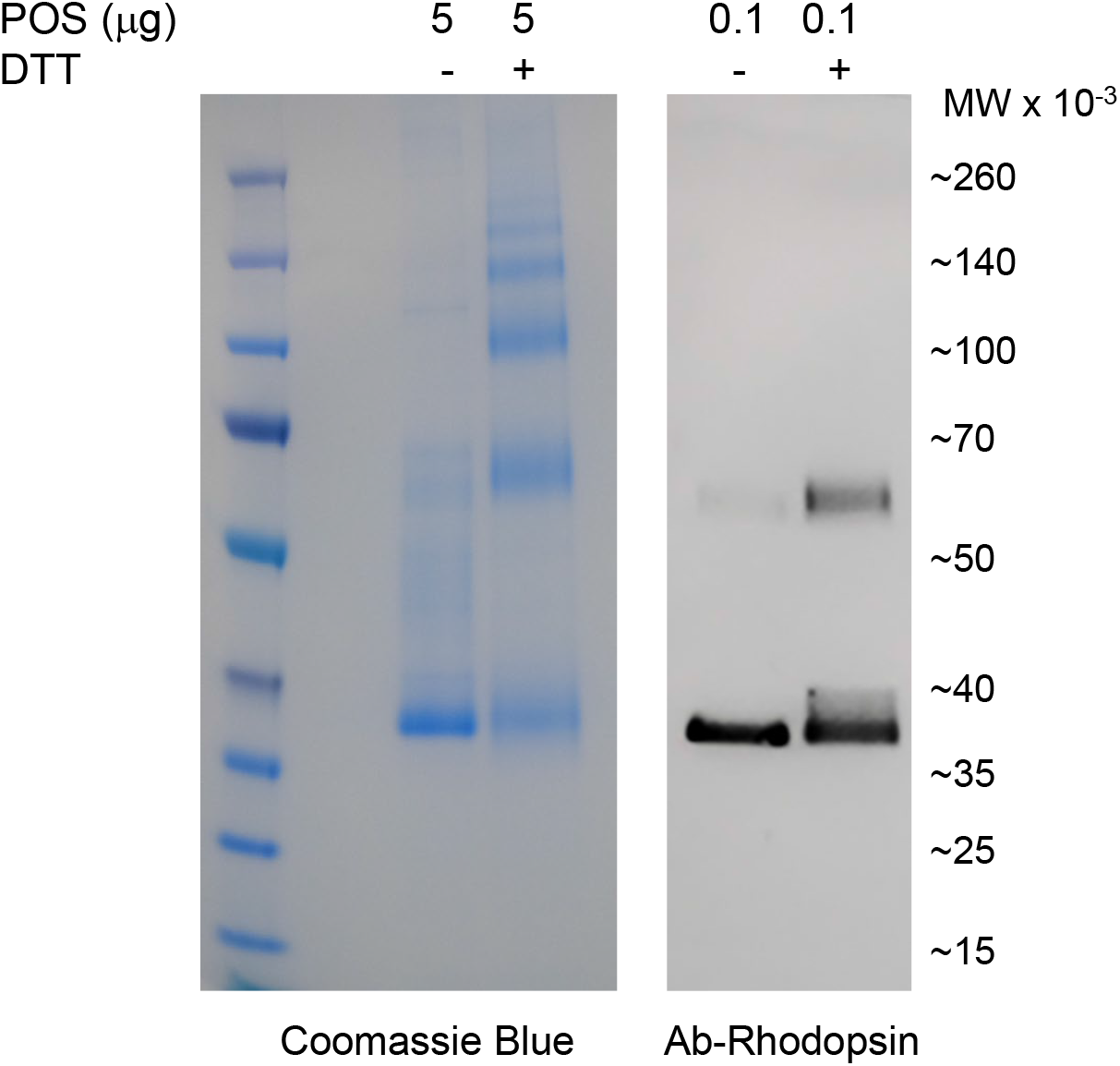
SDS-PAGE and western blot of bovine POS. Proteins in the POS samples were determined and resolved by SDS-PAGE in the same gel in two sets: one with 5 μg and another with 0.1 μg protein per lane. For each set, one sample was non-reduced and the other was reduced with DTT. After electrophoresis, the gels were cut in half lengthwise. The gel portion with 5 μg of protein was stained with Coomassie Blue and the other portion with 0.1 μg protein was transferred to a nitrocellulose membrane for immunostaining using anti-rhodopsin antibodies (as described in Methods). Photos of the stained gel and western blot are shown. The proteins of POS isolated from bovine retina had the expected migration pattern for both reduced and non-reduced conditions, and the main bands stained with Coomassie Blue comigrated with rhodopsin-immunoreactive proteins in western blots of POS proteins.

**Figure S2.**
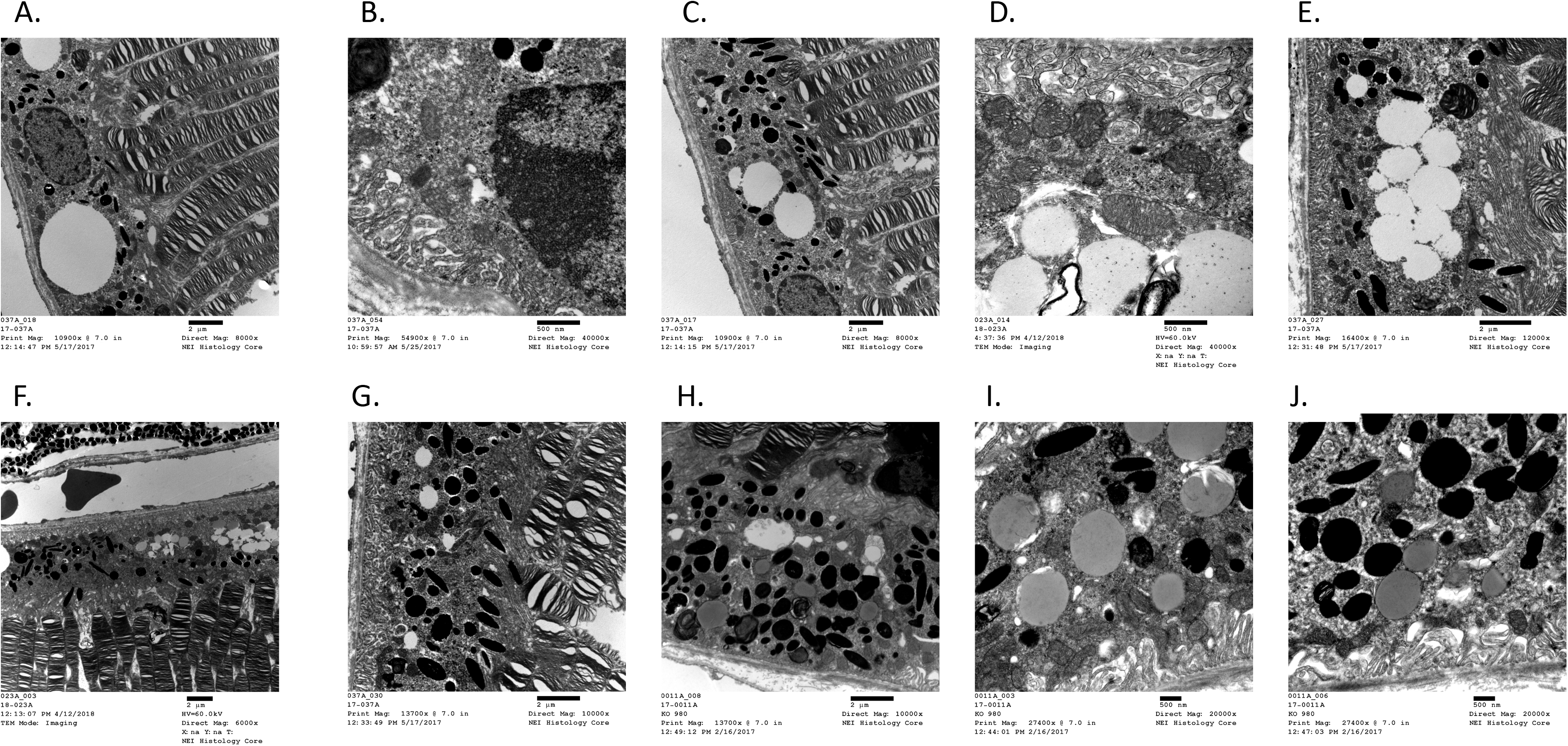
TEM of RPE in RPE-*Pnpla2*-cKO mice. The presence of LDs was associated with lack (Fig. S2A) of or the decreased thickness of the basal infoldings, and with granular cytoplasm, abnormal mitochondria (Fig. S2B), and disorganized localization of organelles (mitochondria and melanosomes) (Fig. S1A). In some cells, the large LDs crowded the cytoplasm and clustered together the mitochondria and melanosomes into the apical region of the cells (Figs. S2A, S2C, S2D); however, LDs number and expansion within the cells appeared to be random and their expansion could go into any direction (Fig. S2E). Normal apical cytoplasmic processes were lacking; however, degeneration in the outer segment (OS) tips of the photoreceptors was visible (Figs. S2A, S2F);. Additionally, normal phagocytosis of the OS was lacking indicating an impaired RPE phagocytosis (Figs. S2A, S2E, S2G). There were apparent unhealthy nuclei with pyknotic chromatin and leakage of extranuclear DNA (enDNA), indicating that the beginning of the necrotic process had started (Fig. S2B). Some RPE cells lacked basal infoldings, normally seen at the basal side (Fig. S2H). Occasionally some RPE cells had lighter low-density cytoplasm indicating degeneration of cytoplasmic components in contrast to the denser and fuller cytoplasm in the RPE of the littermate control (Fig. S2I, S2J).

**Figure S3.**
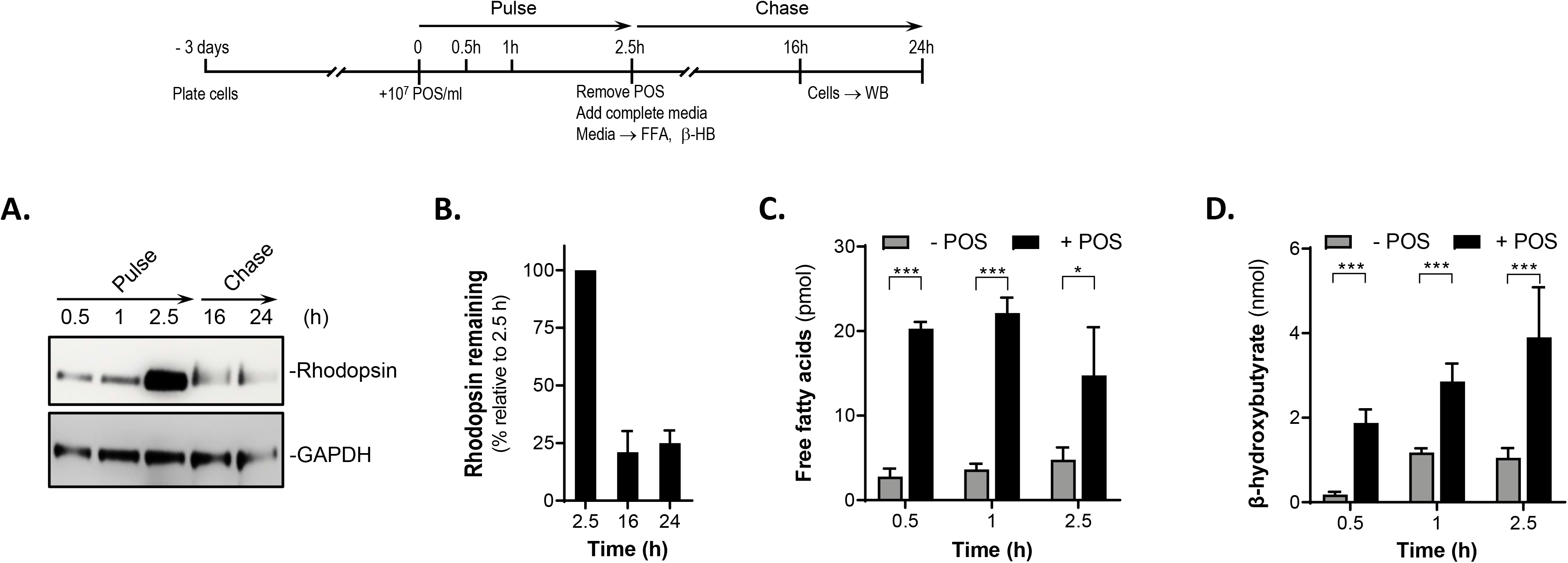
Phagocytosis in ARPE-19 cells. ARPE-19 cells were cultured in 24-well plates for 3 days, and then exposed to POS at 1×107 units/ml for up to a 2.5-h pulse followed by an upto 24-h chase period as described in Methods. (A) Representative immunoblots of total cell lysates during pulse-chase (times indicated at the top of the blot) with anti-rhodopsin followed by reprobing with anti-GAPDH as the loading control are shown. Migration positions of rhodopsin and GAPDH are indicated to the right of the blot. Duplicate biological replicates were performed. (B) Quantification of rhodopsin from duplicate samples per condition from pulse-chase experiments at time periods indicated in the x-axis as from panel (A). Intensities of the immunoreactive bands from duplicate samples of cell lysates were determined. The percentage of the remaining rhodopsin after 16-h chase relative to rhodopsin at 2.5 h-pulse was plotted. (C-D) Levels of free fatty acids (C) and ⍰-HB (D) measured in culture media of cells incubated with and without POS for the indicated periods of time (x-axis) were plotted and shown. n = 3 Data are presented as means ± S.D. * p < 0.05, ***p<0.001.

**Figure S4.**
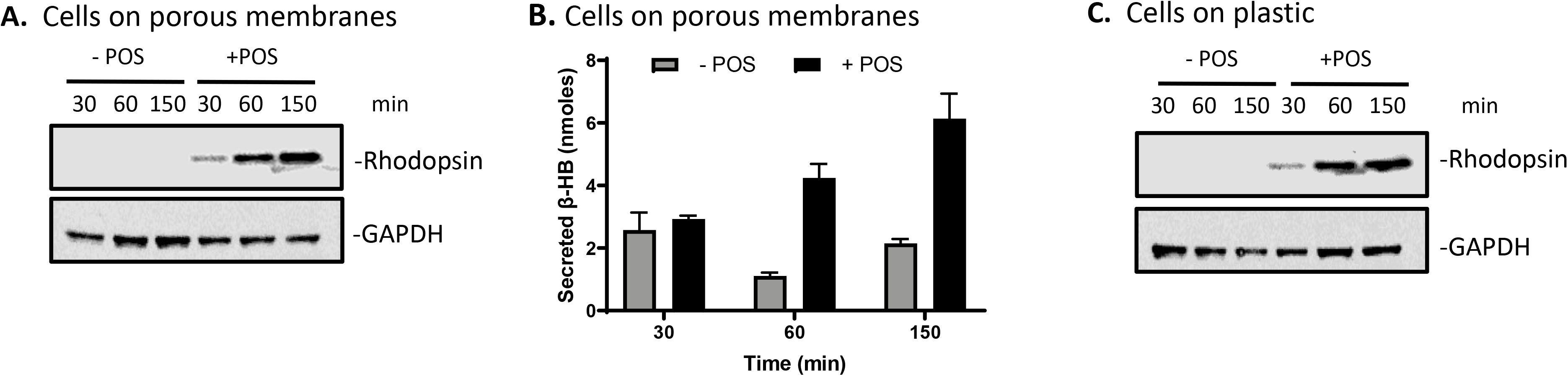
Phagocytosis in ARPE-19 cells in porous membranes. ARPE-19 cells were treated with 1×107 POS/ml. (A) Representative immunoblot showing rhodopsin internalization from total cell lysates of ARPE-19 cells following 30, 60, and 150 min of POS incubation following plating in 12-well transwell inserts for 3 weeks. Cell extracts were resolved by SDS-PAGE followed by immunoblotting with anti-rhodopsin. The blot was stripped and reprobed with anti-GAPDH as a loading control. (B) Levels of B-HB secreted towards the apical membrane of ARPE-19 cells following POS incubation for 30, 60, and 150 min. Data are presented as means ± S.D. **ARPE-19 cells plated on porous membranes engulf bovine outer segments** To demonstrate a functional assay to study phagocytosis in ARPE-19 cells we perform the assay with confluent cells attached on porous membranes **Methods:** ARPE-19 cells seeded on porous membranes were incubated for 3 weeks in culturing media. Then the media was replaced with Ringer’s solution alone or Ringer’s solution containing 1 x 107 POS/ml and 5 mM glucose for the indicated time points. Rhodopsin was detected by western blotting. Rhodopsin levels in the lysates of cells incubated with POS were detected in as little as 30 min and up to 2.5 h following POS incubation, while rhodopsin was undetectable in cells without POS (**Fig. S4A**). B-HB levels released into the media of the apical chamber of transwells following POS incubation increased four-fold and three-fold after 60 and 150 min, respectively, while released B-HB levels from cells incubated with Ringer’s solution alone did not increase (**Fig. S4B**).

**Figure S5.**
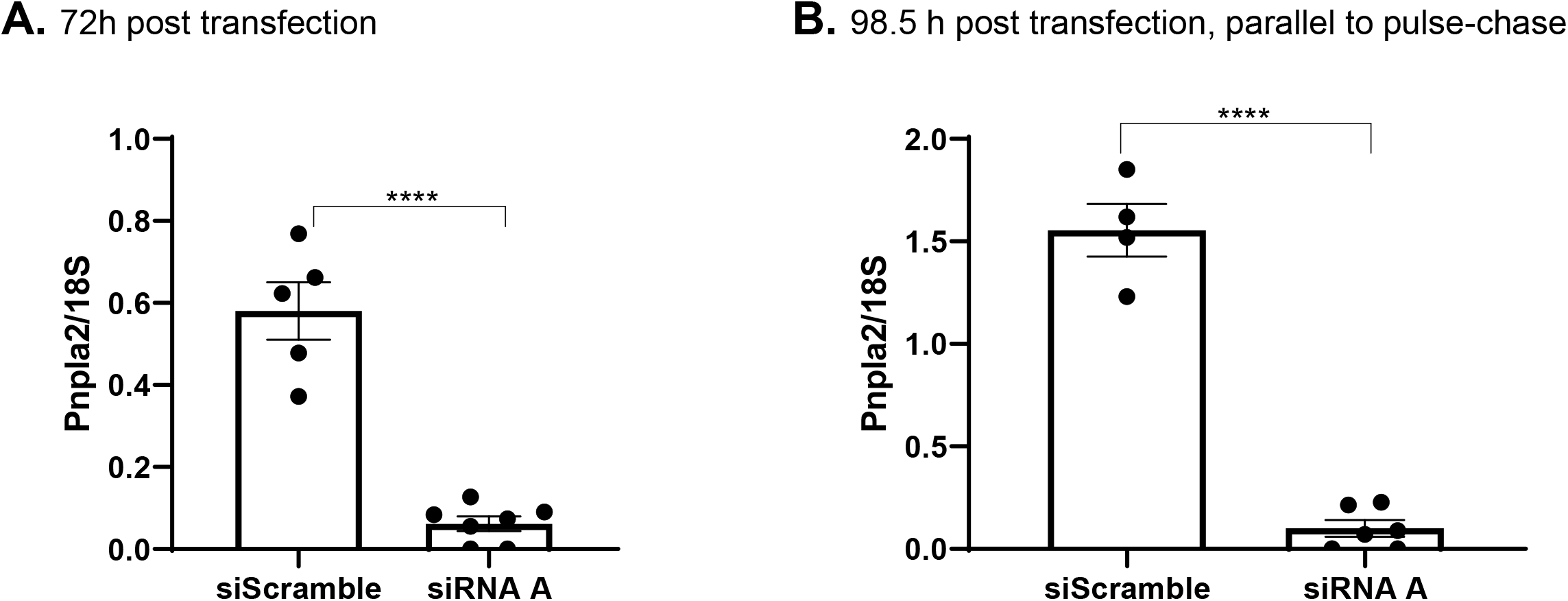
ARPE-19 cells were transfected with siScramble siRNA control or siRNAs targeting *PNPLA2 (siPNPLA2 A)*. RT-qPCR to measure *PNPLA2* mRNA levels in ARPE-19 cells at **(A)** 72 h post-transfection and (**B**) 98.5h post transfection equivalent to pulse (2.5h) and chase (24h) was performed with siRNA duplexes (as indicated in the *x*-axis). Treatment of cells in panel B was as for pulse-chase (see diagram in Fig S3). *PNPLA2* mRNA levels were normalized to 18S. n =3 biological replicates, each data point corresponds to the average of triplicate PCR reactions. The RT-PCR was repeated twice per biological replicate. Values that fell out of the standard curve were not included in the plot. The data shows that si*PNPLA2* duplex silenced *PNPLA2* in ARPE-19 at 72 h post-transfection and that silencing was maintained throughout a 2.5 h and pulse-chase of 24 h.

